# A nitrite-responsive regulatory RNA locus sustains commensal resilience against nitrosative stress

**DOI:** 10.64898/2026.07.09.736866

**Authors:** Ryan T. Fansler, Daniel W. Bak, Madison Langford-Butler, Liang Chen, Deepanshu Singla, Luisella Spiga, Jonathan Livny, John Karijolich, Chance Meers, Qiangjun Zhou, Eranthie Weerapana, Wenhan Zhu

## Abstract

Commensal microbes in the gastrointestinal tract are central to host health, yet they must adapt to frequent perturbations such as intestinal inflammation that challenges microbial homeostasis. A major challenge during inflammation is exposure to host-derived reactive nitrogen species (RNS), which damage macromolecules and impair microbial fitness, but how commensals orchestrate defense against nitrosative stress remains poorly defined. Here, we show that *Bacteroides thetaiotaomicron* mounts a protective RNS-defense program centered on the hybrid cluster protein Hcp, which is required for fitness under nitrosative stress. We identify a nitrite-responsive SnoA locus (Stress-responsive Nitric Oxide regulator A) that promotes HcpR-dependent *hcp* expression. *In vivo*, this pathway promotes commensal resilience in both an antibiotic-perturbed, Nos2-dependent model of intestinal nitrosative stress and during *Salmonella*-induced gut inflammation. Together, our findings identify a regulatory pathway that enables a dominant gut commensal to withstand host-derived nitrosative stress and persist during intestinal inflammation.

## Introduction

Inflammation radically alters the chemical landscape of the gut. As activated immune cells infiltrate the mucosa, the host deploys a powerful arsenal of antimicrobial strategies, including the production of copious reactive nitrogen species (RNS), such as nitric oxide (NO)^1^. NO is a small, diffusible radical with broad reactivity toward metals and thiol-containing macromolecules. Its reactions disrupt enzymes that mediate central metabolism, redox balance, and nucleotide biosynthesis, leading to impaired microbial fitness^1,2^. As NO diffuses outward from inflamed tissues, it reacts with reactive oxygen species to form peroxynitrite (ONOO^−^), which can further decompose or react to generate nitrite (NO_2_^−^) and nitrate (NO_3_^−^)^3^. Although both molecules can serve as respiratory electron acceptors, nitrate primarily functions as a microbial respiratory substrate, whereas nitrite remains highly reactive and exerts potent antimicrobial effects^4^. Moreover, nitrite is relatively stable in the intestinal lumen and can be converted back into reactive nitrogen species under appropriate physiological conditions, thereby prolonging nitrosative stress^5^. Consequently, exposure of bacterial cells to NO and NO_2_^−^ results in periods of growth arrest^6^. These chemical defenses substantially constrain pathogen expansion^7^, but they also impose significant collateral stress on resident commensals. Yet, the molecular targets and protective strategies that allow gut commensals to endure nitrosative stress remain poorly understood.

In contrast to well-characterized pathogens, which wield an impressive array of RNS detoxification systems^1^, how commensals withstand and survive these attacks remains largely undefined. However, their survival under inflammatory stress is central to gut health, as these microbial residents are vital for maintaining metabolic homeostasis, modulating the immune system, and providing colonization resistance against pathogens^8,9^. Inflammatory nitrosative bursts can deplete key commensals, reshaping microbiome composition and function,^10^ and exacerbating chronic conditions such as infectious diarrhea and inflammatory bowel disease^11^. Thus, elucidating how health-associated commensals sense and withstand RNS is essential for elucidating how the human microbiome preserves its stable community structure and function over time^12^.

Work in model pathogens has defined a suite of enzymatic defenses that detoxify RNS, including the NO-detoxifying flavoenzymes Hmp and NorV and iron-dependent nitrite-reducing systems, such as NirB and NrfA^1^. More recently, the hybrid cluster protein Hcp has emerged as a high-affinity NO reductase with an important role in mitigating nitrosative stress^13^. In parallel, bacteria can buffer RNS toxicity by remodeling metabolism to bypass damaged biosynthetic reactions,^14^ increase uptake or salvage of essential metabolites, and redirect flux through alternative pathways that compensate for nitrosative damage^15^. Bacteria can also replace labile metals such as Fe with more RNS-resistant cofactors such as Mn^16–18^. These diverse nitrosative defenses are regulated by a wide range of transcription factors, including NorR, NsrR, and HcpR, which respectively sense nitrosative stress via direct NO binding, iron–sulfur cluster damage, or heme interaction^19–22^, and subsequently bind to the promoter regions of the RNS-defense effector genes. Beyond transcription factors, regulatory non-coding RNAs, including small RNAs (sRNAs)^23^, have recently been implicated in regulating a broad range of bacterial stress responses^24^, including emerging roles in nitrosative stress tolerance^25,26^. However, the mechanisms by which regulatory RNAs coordinate bacterial defenses against nitrosative stress remain largely undefined.

Here, we investigate how the prominent gut commensal *Bacteroides thetaiotaomicron* withstands nitrosative stress. We define protein targets damaged by RNS and show that *B. thetaiotaomicron* mounts a coordinated transcriptional response that relies on NrfA and Hcp to maintain fitness *in vitro* and during intestinal inflammation. We further identify a nitrite-responsive SnoA regulatory RNA locus that promotes HcpR-dependent activation of *hcp* and contributes to commensal fitness under nitrosative stress. Together, our findings uncover a nitrosative-defense strategy that preserves commensal resilience during inflammation and reveal an RNA-associated regulatory mechanism controlling bacterial stress adaptation.

## Results

### Nitrosative and iron stresses synergistically impair commensal fitness by targeting central metabolism and biosynthetic pathways

To define how a gut commensal withstands nitrosative stress, we exposed *B. thetaiotaomicron* cultures to increasing concentrations of the nitric oxide donor DPTA NONOate^27^ or nitrite. In rich medium, *B. thetaiotaomicron* tolerated both RNS well, maintaining robust growth even at high micromolar to low millimolar doses (**Fig. 1A&B**). During intestinal inflammation, however, nitrosative stress is invariably coupled to nutritional immunity^28,29^, which restricts the bioavailability of metals, including iron, a cofactor required for many canonical RNS detoxification systems^30^. We therefore repeated these experiments in the presence of the iron chelator bathophenanthrolinedisulfonate (BPS)^31^, which models inflammation-associated iron limitation. Under these conditions, both NO and nitrite caused pronounced growth defects (**Fig. 1C&D**), indicating that nitrosative and iron stresses synergize to create a substantial challenge for *B. thetaiotaomicron*.

**Fig. 1.**
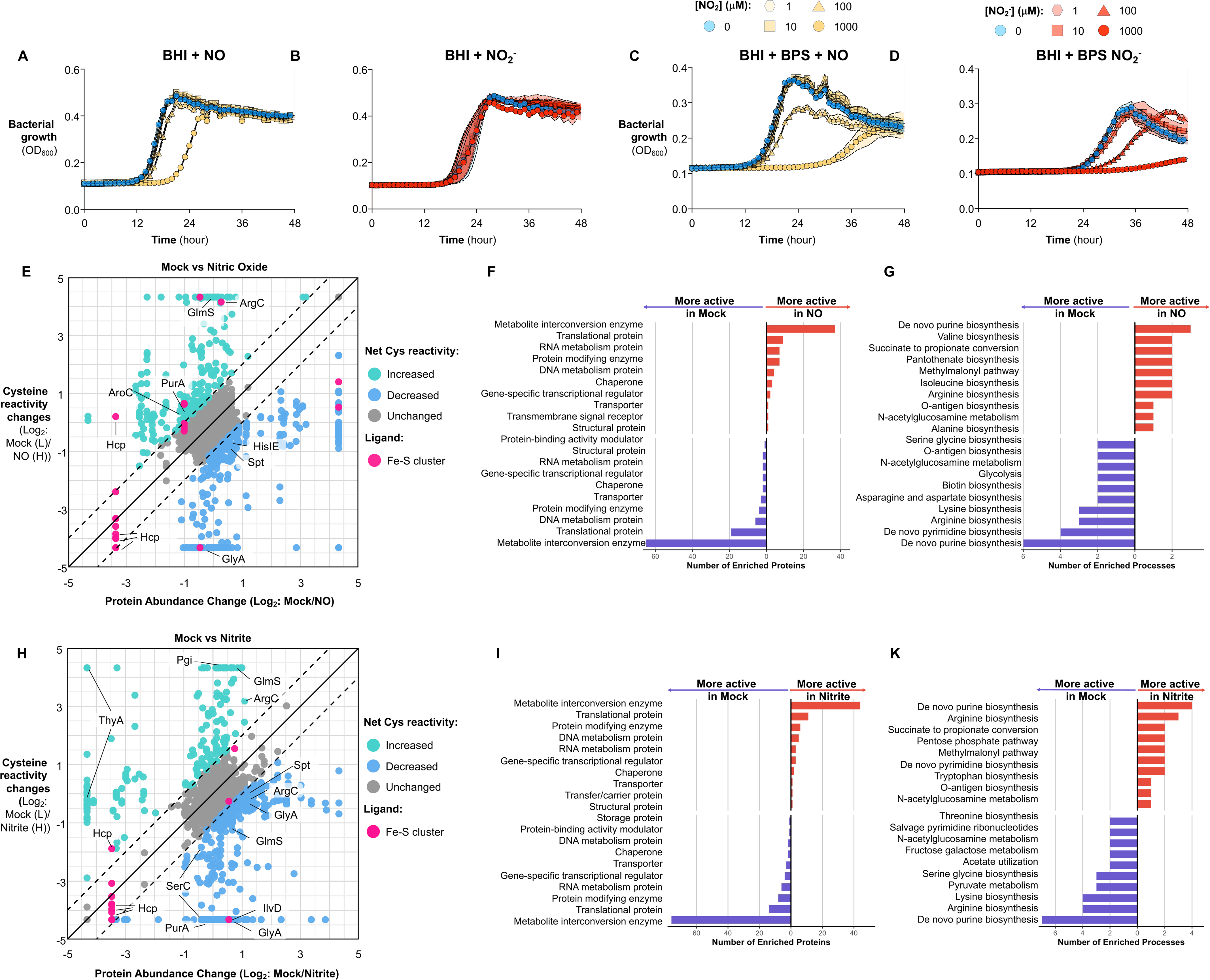
Nitrosative stress impairs commensal fitness and remodels cysteine reactivity. **A-D.** Growth of *B. thetaiotaomicron* in BHI (Brain Heart Infusion) under iron-limiting conditions (BPS) with the indicated nitrosative stressors. Bacterial growth (OD_600_) was monitored over time. **A-B,** nitric oxide (NO) and nitrite under iron replete condition; **C-D,** nitric oxide (NO) and nitrite under iron limited condition (BPS). **E-K.** Quantitative cysteine chemoproteomics (isoTOP-ABPP using IA-alkyne) after exposure to nitric oxide (**E-G**) or nitrite (**H-K**). **E, H,** two-dimensional datasets plotting cysteine probe labeling under control vs. stress: points above the unity/expected line indicate increased cysteine reactivity, on the line indicate no net change, and below the line indicate decreased reactivity. **F, I,** PANTHER protein-class enrichment among sites with altered cysteine activity. **G, K,** PANTHER biological process enrichment for the same sets. **E&H:** ArgC (N-acetyl-γ-glutamyl-phosphate reductase), AroC (chorismate synthase), GlyA (serine hydroxymethyltransferase), GlmS (glutamine—fructose-6-phosphate aminotransferase), Hcp (hybrid cluster protein), HisIE (Histidine biosynthesis bifunctional protein), IlvD (dihydroxy-acid dehydratase), Pgi (glucose-6-phosphate isomerase), PurA (adenylosuccinate synthetase), SerC (phosphoserine aminotransferase), Spt (serine palmitoyltransferase), and ThyA (thymidylate synthase).

We next investigated how nitrosative stress perturbs commensal physiology. We focused on the impact of nitrosative stress on the proteome-wide reactivity of cysteine residues, as they are key redox-sensitive nucleophiles that both coordinate metal cofactors and support catalysis across a vast array of cellular functions^32^. Reactive nitrogen species disrupt cellular physiology by targeting cysteine-centered chemistry, including direct modification of reactive cysteine thiols and damage to cysteine-coordinated iron-sulfur clusters^6^. Cysteine nitrosation can alter protein function and signaling, whereas NO-mediated degradation of iron-sulfur cofactors disables enzymes involved in respiration, metabolism, and stress resistance^33,34^. Although these mechanisms are well established in pathogens, how RNS globally remodel cysteine reactivity across commensal proteomes remains largely unknown.

To globally profile nitrosative stress sensitivity in *B. thetaiotaomicron*, we applied a modified isotopic tandem orthogonal proteolysis-activity-based protein profiling (isoTOP-ABPP) platform^35^. This method employs the cysteine-reactive probe iodoacetamide-alkyne (IA-alkyne) to quantify changes in cysteine reactivity proteome-wide during nitrosative stress. *B. thetaiotaomicron* cultures were treated with the NO donor DPTA NONOate, nitrite, or vehicle, differentially labeled with light (vehicle) or heavy (NO or nitrite) IA-alkyne, and then analyzed by LC-MS/MS. Probe-derived cysteine reactivity log_2_(L/H) ratios were then normalized to protein abundance, and the resulting net cysteine log_2_(L/H) ratios (*R*_C_) report relative changes in IA-alkyne labeling, with *R*_C_ < −1 indicating cysteine reactivity under nitrosative stress (e.g., nitrosation of Fe-S clusters that exposes coordinating cysteines) and *R*_C_ > 1 indicating decreased cysteine reactivity (e.g., direct cysteine nitrosation that blocks IA-alkyne probe access) (**Fig. S1A**). Both NO and nitrite broadly remodeled cysteine reactivity across the proteome, with NO altering 249 of 890 quantified cysteines and nitrite altering 265 of 953 quantified cysteines (∼28% for each condition) (**Fig. S1B,C**; **Fig. 1E,H, Supplementary Table S1**). Proteins enriched with nitrosative-sensitive cysteine residues mapped to biological pathways involved in nucleic acid metabolism, central carbon metabolism, and amino-acid biosynthetic pathways **(Fig. 1F, G, I, K**).

Consistent with damage to these pathways underlying *B. thetaiotaomicron’s* growth impairment during nitrosative stress, supplementation with casamino acids, pyruvate, or glutamate significantly rescued the bacterium’s growth in a semi-defined medium challenged with nitrite (**Fig. S1D-H**). Supplementation of the amino acid glutamate alone was tested due to glutamate synthase and glutamate dehydrogenase being top hits in both the nitrite and nitric oxide chemoproteomic screens. Notably, supplementation with glutamate alone rescued growth to a comparable degree as the consortium of casamino acids (**Fig. S1I**), suggesting that glutamate biosynthesis constitutes a key vulnerability for *B. thetaiotaomicron* during nitrosative stress. Indeed, *B. thetaiotaomicron* cultures challenged with nitric oxide displayed markedly lower intracellular glutamate levels than those of mock-treated controls (**Fig. S1J**).

### *B. thetaiotaomicron* mounts a protective transcription program to defend against nitrosative stress

The observed tolerance of *B. thetaiotaomicron* to stress suggests that this commensal deploys dedicated defense mechanisms (**Fig. 1**). To define these pathways, we profiled genome-wide transcriptional changes following exposure to nitrite, an experimentally tractable RNS^36^ that phenocopies many of the proteomic and physiological consequences of NO, including widespread remodeling of cysteine reactivity and impaired bacterial fitness (**Fig. 1**). As expected, *B. thetaiotaomicron* mounted a robust transcription response to nitrite (**Fig. 2A&S2A**), differentially regulating genes involved in central metabolism, nucleotide and vitamin biosynthesis, as well as nitrosative stress defense pathways (**Fig. 2B&C**). Among the most strongly upregulated genes were hybrid cluster protein (*hcp*) and cytochrome c nitrite reductase (*nrfA*, hereafter referred to as NR), two conserved components of bacterial nitrosative-stress defense systems^37,38^. In contrast, nitrate triggered a distinct redox-associated response (**Fig. S2B-F**) despite not impairing *B. thetaiotaomicron* growth (**Fig. S2G-H**). Curiously, nitrate also induced *hcp* expression despite its overall lack of toxicity (**Fig. S2C**), suggesting that either *B. thetaiotaomicron* generates low background levels of nitrite from nitrate under anaerobic conditions or nitrate may perturb intracellular redox and Fe-S homeostasis, the primary signals for transcription regulators that control *hcp* expression in strict anaerobes^20,39,40^.

**Fig. 2.**
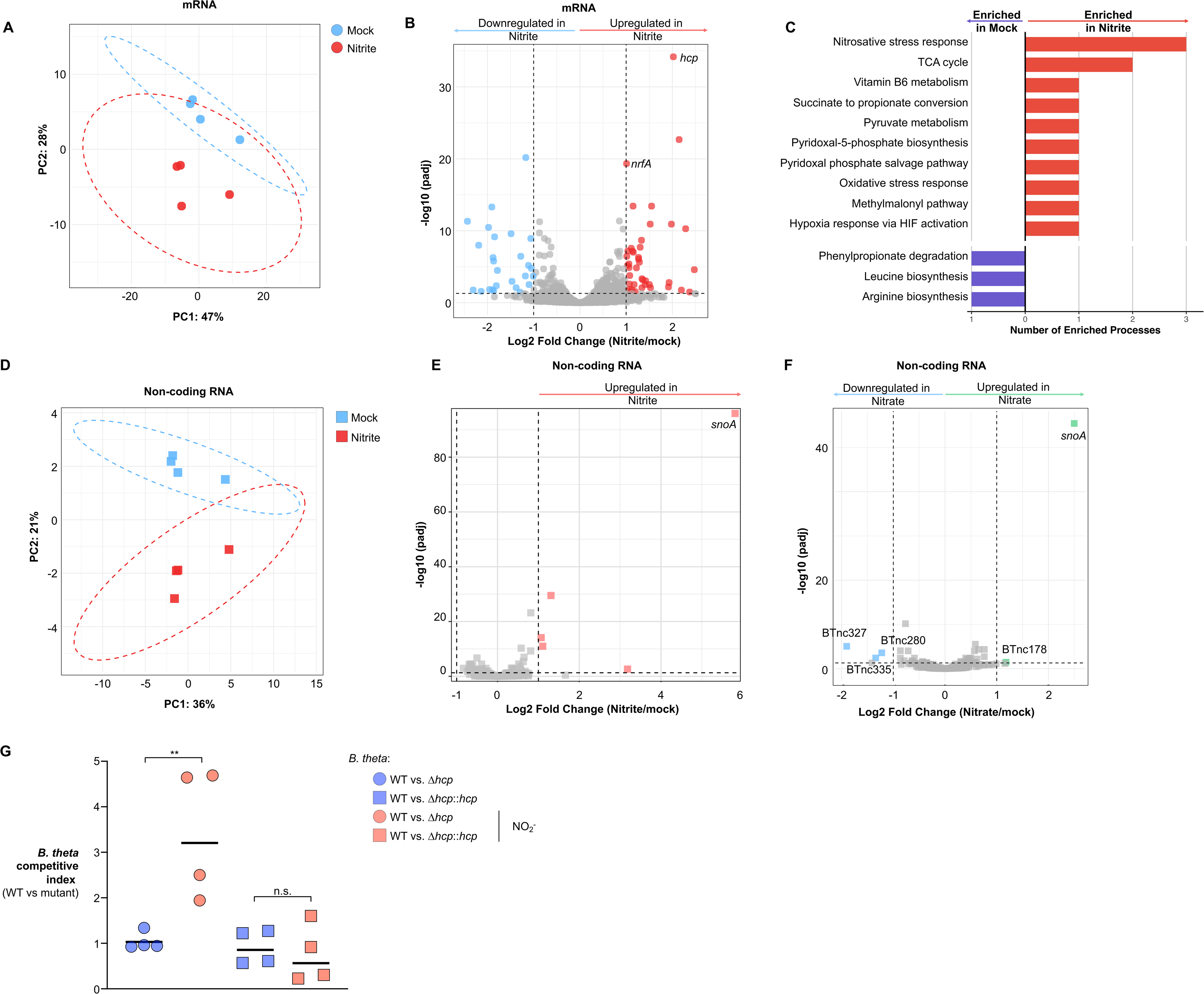
*B. thetaiotaomicron* mounts a protective transcriptional program to detoxify nitrosative stress. **A-F.** *B. thetaiotaomicron* cultures were exposed to nitrite (**A-E**) or nitrate control (**F**) and transcriptome profiled by RNA-seq. **A,** principal component analysis (PCA) of the mRNA transcriptome. **B,** mRNA volcano plot. **C,** PANTHER biological-process enrichment of differentially expressed mRNAs. **D,** PCA and **E,** volcano plot of non-coding RNA transcriptome upon nitrite exposure. **F,** non-coding RNA volcano plot of *B. thetaiotaomicron* cultures exposed to nitrate control. **G.** A 1:1 mixture of *B. thetaiotaomicron* wild-type and an isogenic *hcp*-deficient mutant was inoculated in BHI with iron limitation (BPS) ± nitrosative stress (NO_2_^−^ or NO). Competitive index was determined by selective plating. Bars (where shown) depict geometric means. *, *P* < 0.05; **, *P* < 0.01. BPS, bathophenanthroline disulfonate; NO_2_^−^, nitrite; DPTA NONOate, Dipropylenetriamine NONOate (nitric oxide producer).

In addition to mRNA, we evaluated changes in the annotated non-coding RNA fraction of the transcriptome, as regulatory RNAs have been implicated in diverse bacterial stress responses, including nitrosative stress tolerance^24,41,42^. Nitrite elicited a similarly robust non-coding RNA response (**Fig. 2D-E**), with BTnc284, a previously reported non-coding RNA^41^, representing the most strongly induced annotated non-coding RNA feature after nitrite exposure. Although previous transcriptome analyses annotated this region as BTnc284, the precise structure of the mature RNA remains unresolved. Throughout this study, we therefore refer to the transcriptionally active regulatory region as the SnoA locus (Stress-responsive Nitric Oxide regulator A). Notably, nitrate also induced the SnoA locus (**Fig. 2F, S2F**), consistent with the hypothesis that nitrate perturbs redox or Fe-S homeostasis rather than directly causing nitrosative stress^39,40,43^.

Because *nrfA* (NR) activity protects *B. thetaiotaomicron* only under iron-limiting conditions (**Fig. S2I**), we reasoned that *hcp*, which encodes a hybrid-cluster protein containing an unusual Fe/S/O metallocluster^38^, may provide a complementary detoxification pathway. Although Hcp was initially described as a high-affinity NO reductase in *E. coli*^38,44^, accumulating evidence shows that Hcps broadly function as nitrosative-stress defense enzymes across anaerobes^45,46^. To test whether Hcp contributes to *B. thetaiotaomicron* nitrosative resilience, we constructed a non-polar deletion mutant of *hcp* and performed a competition assay under nitrosative stress and iron-limiting conditions. Consistent with our hypothesis, wild-type *B. thetaiotaomicron* significantly outcompeted the Δ*hcp* mutant when challenged with nitrite, even under iron-replete conditions (**Fig. 2G**). This fitness defect was rescued by complementation of *hcp* at a neutral genomic locus. Moreover, upon nitrite challenge under iron-replete conditions, loss of *hcp*, but not loss of NR, led to elevated residual nitrite in spent medium, and the ΔNR Δ*hcp* double mutant phenocopied the Δ*hcp* single mutant (**Fig. S2J**). Together, these findings indicate that B. thetaiotaomicron uses complementary nitrosative-stress defenses, with Hcp serving as the dominant contributor under iron-replete nitrite stress and NrfA providing additional protection under iron-limited conditions.

### The SnoA regulatory RNA locus modulates nitrosative-stress defense in *B. thetaiotaomicron*

In our transcriptome data, we found that the SnoA-associated non-coding RNA feature was highly upregulated in response to nitrite exposure (**Fig. 2E**), suggesting a role in nitrosative-stress defense in *B. thetaiotaomicron*. Intriguingly, *snoA* is encoded within the promoter region of *hcp* but in the opposite orientation, downstream of the transcription factor HcpR, which has been implicated in licensing *hcp* transcription in response to nitrosative stress^46,47^. A previous high-resolution survey of the *B. thetaiotaomicron* transcriptome^41^ annotated the SnoA region and placed the predicted transcript opposite the −7 element^48^ of the hcp promoter (**Fig. 3A**). This arrangement suggested a possible interaction between RNA arising from the SnoA locus and the *hcp* promoter (P*_hcp_*), motivating us to investigate its regulatory function.

**Fig. 3.**
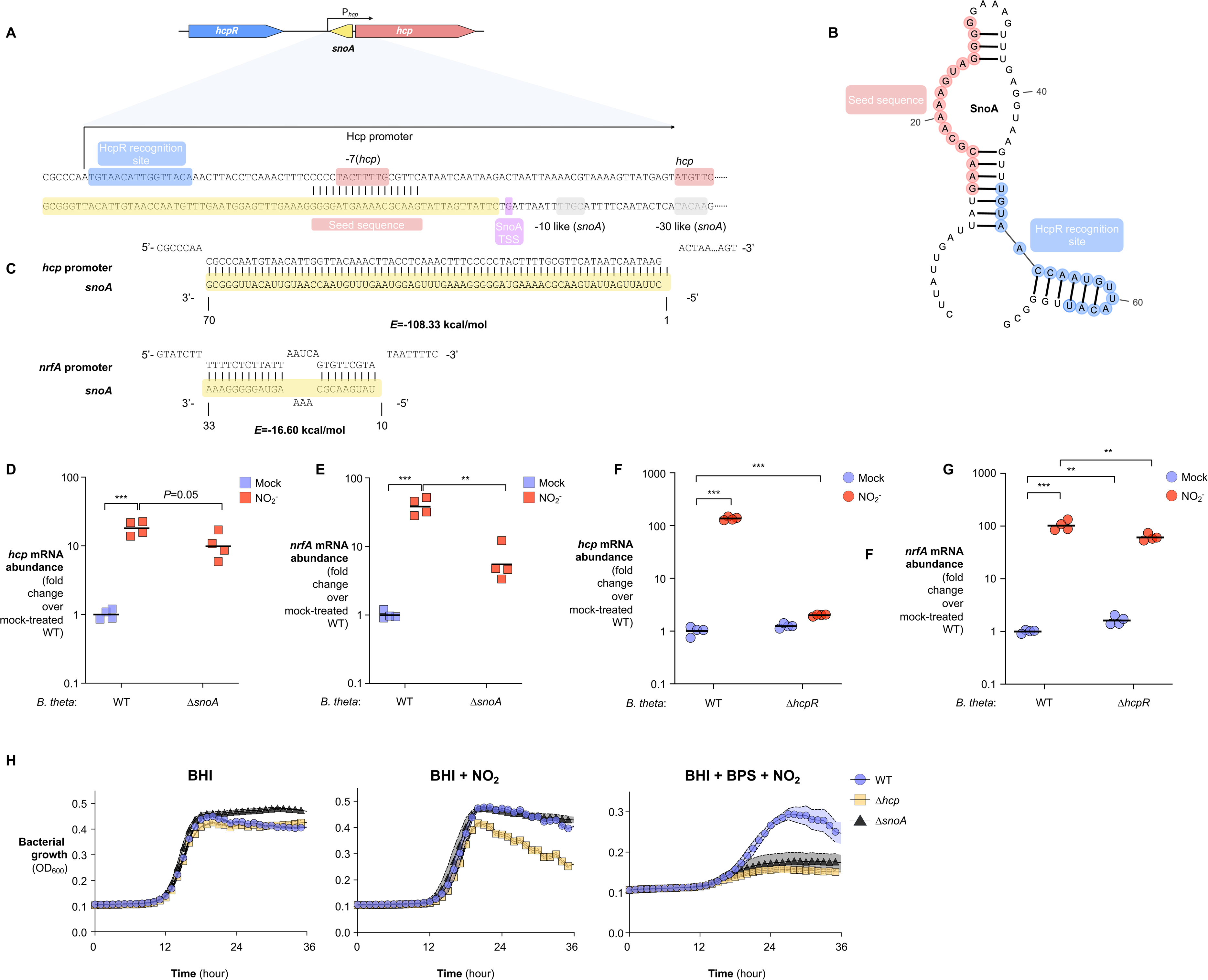
The SnoA regulatory RNA locus modulates nitrosative-stress defense in *B. thetaiotaomicron*. **A.** Genomic locus schematic showing *hcpR* (transcription factor), *snoA* locus, and *hcp* (hybrid cluster protein). **B.** Predicted minimum-free-energy secondary structure (hairpin) of SnoA. **C.** IntaRNA predictions of SnoA base-pairing with the promoter regions of *hcp* and *nrfA*. **D-G.** RT-qPCR of *hcp* (**D, F**) and *nrfA* (**E, G**) mRNA in the indicated strains after exposure to nitrite or mock control. **H.** Growth of *B. thetaiotaomicron* cultures (OD_600_) under iron limitation with nitrosative stress. Bars show geometric means. **, *P* < 0.01; ***, *P* < 0.001.

Aligning SnoA with putative homologues from other *Bacteroides* species using CopraRNA^49^ revealed a highly conserved segment that could function as a seed region involved in base-pairing between SnoA and P*_hcp_* (**Fig. S3A**). Notably, this putative seed sequence is predicted to remain unpaired within SnoA itself^50^ and overlaps precisely with the −7 element of the *hcp* promoter (**Fig. 3A&B**). Consistently, IntaRNA predicts that RNA arising from the SnoA locus binds to the *hcp* promoter with high affinity (*E* = - 108.33 kcal/mol) and also binds the *nrfA* promoter with a moderate affinity (*E* = −16.60 kcal/mol, **Fig. 3C**), suggesting that RNA arising from the SnoA locus may contribute to coordinated regulation of *hcp* and *nrfA*.

To test this, we sought to disrupt *snoA* without abolishing *hcp* induction. Because *snoA* is encoded antisense to the *hcp* promoter, we first replaced the entire *hcp* promoter with an alternative promoter that is responsive to nitrosative stress (BT_1143, L2FC = 2.14; *hcp*, L2FC = 2.02; mock vs. nitrite, **Supplementary Table S2**). This strategy allowed us to generate a Δ*snoA* strain in which *hcp* remains nitrite-inducible but is no longer under SnoA control. Indeed, *hcp* transcription in the Δ*snoA* strain remained responsive to nitrite, although the magnitude of induction was reduced compared to the wild-type strain (**Fig. 3D**), supporting a positive regulatory contribution from the SnoA locus. In line with this model, *snoA* deletion caused a significant reduction in *nrfA* transcription (**Fig. 3E**), consistent with the partial complementarity between SnoA and *nrfA* promoter and a role for the SnoA locus in supporting full *nrfA* induction (**Fig. 3C**).

Curiously, deletion of *hcpR*, the transcription factor located upstream of *snoA* and *hcp*, completely abolished *hcp* transcription but had only a modest, albeit significant, impact on *nrfA* expression and no effect on *snoA* levels (**Fig. 3F-G, S3B**). These data indicate that SnoA and HcpR are both positive regulators of the *B. thetaiotaomicron* nitrosative-stress response, but they act through distinct mechanisms and targets. Consistent with this notion, mutants lacking *hcp* or *hcpR* exhibited fitness defects in the presence of nitrosative stress, in the form of 50 µM sodium nitrite, which were more pronounced under iron-limited conditions (**Fig. 3H, S3C**). Notably, the Δ*hcp* mutant strain displayed almost complete growth arrest at higher levels of nitrosative stress (1 mM nitrite), even in iron-replete media (**Fig. S3D**). By contrast, the Δ*snoA* mutant showed measurable growth defects only when both nitrosative and iron stresses were present, likely because *hcp* and *nrfA* can still be partially induced in the absence of SnoA (**Fig. 3H, S3D**). A strain lacking both *snoA* and *hcp* also displayed strong fitness impairment under combined iron and nitrosative stress, and these defects were partially rescued by reinserting the *snoA*-*hcp* locus into a neutral site in the genome (**Fig. S3C&E**). Together, these data indicate that HcpR and SnoA cooperate to control *hcp* transcription in response to nitrosative stress, and that additional factors likely contribute to full *nrfA* induction beyond SnoA alone.

### SnoA-associated RNA can form an HcpR-containing complex with target promoter DNA

In bacteria, transcription factors typically regulate gene expression by sensing environmental cues, binding specific DNA motifs, and interacting with RNA polymerase at target promoters^51,52^. Consistent with this paradigm, the *B. thetaiotaomicron* HcpR binds heme, a physiologic indicator of nitrosative stress^53^ (**Fig. S4A**). Accordingly, induction of *hcp* expression by challenge with nitrite is highly iron-dependent, as the induction of *hcp*, but not *nrfA,* was significantly ablated when *B. thetaiotaomicron* was challenged with nitrite and iron limitation simultaneously (**Fig. S4B**). This observation is consistent with *hcp* expression’s stronger reliance on the heme-binding HcpR (**Fig. 3F-G**). However, the perfect complementarity between the SnoA-associated RNA sequence and the *hcp* promoter suggested a distinct regulatory mode in which RNA arising from this locus may participate in HcpR-dependent *hcp* transcription.

Alignment of SnoA with its homologs revealed a well-conserved region (**Fig. S3A**) that pairs precisely with a conserved HcpR recognition sequence within the *hcp* promoter, originally defined in *P. gingivalis*^46,54^ (**Fig. S4C**). Covariance analysis of HcpR and SnoA across bacterial genomes identified two highly conserved sequence motifs, including a 3’ hairpin in SnoA that aligns with the putative HcpR binding site in the *hcp* promoter (**Fig. 4A**). Because RNA hairpins frequently serve as protein-binding modules^55^, and because of the complementarity between the SnoA-associated RNA sequence and the *hcp* promoter, we hypothesized that RNA arising from the SnoA locus contributes to assembly or stabilization of an HcpR-promoter complex. Supporting this, AlphaFold3^56^ predictions indicate that HcpR engages SnoA with a prominent interface at the 3’ hairpin (**Fig. 4B**). Moreover, HcpR, SnoA, and the *hcp* promoter (P*_hcp_*) are predicted to form a ternary complex in which an α-helix of HcpR contacts the DNA:RNA duplex at the HcpR recognition site (Pro187/Ser188/Arg191, **Fig. 4C**). The same helix is also predicted to contact the DNA:RNA duplex at the *nrfA* promoter (**Fig. S4D**). Although the primary sequences of the *hcpR*-*snoA*-*hcp* locus are only moderately conserved across Bacteroides, and SnoA is highly divergent outside the HcpR-recognition and seed regions (**Fig. 4A**), comparative phylogenetic analyses of both Hcp and HcpR homologs identified SnoA-associated loci within distinct clades (**Fig. S4E**). Consistent with this clade-restricted distribution, AlphaFold3 modeling predicted similar HcpR–SnoA–P*_hcp_* architectures across representative species (**Fig. 4D**), suggesting that this RNA-associated regulatory mechanism extends beyond *B. thetaiotaomicron* while remaining restricted to a subset of *Bacteroides* lineages.

**Fig. 4.**
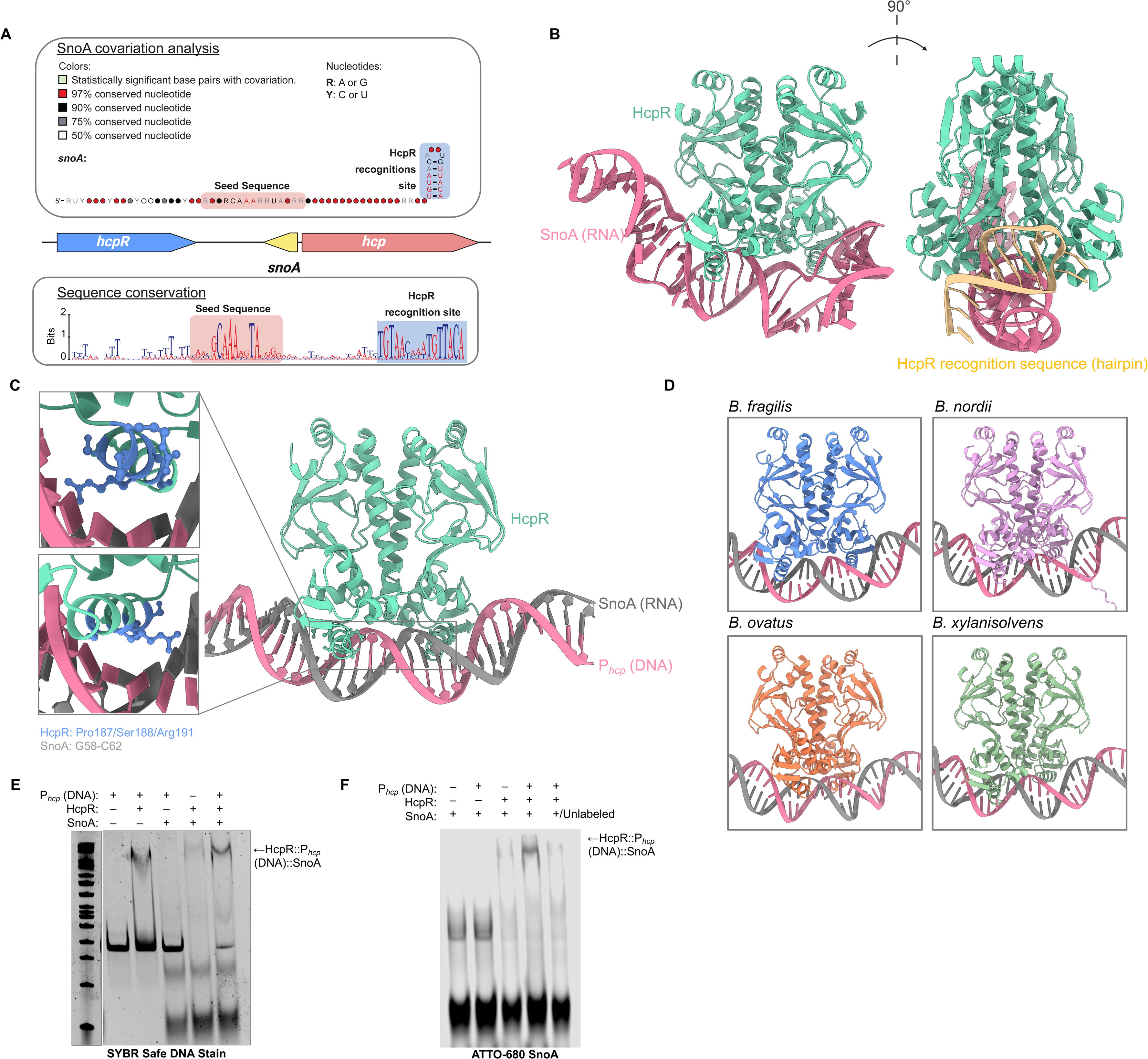
SnoA-associated RNA forms HcpR-containing complexes with conserved promoter elements. **A.** Covariance model (CM) of SnoA highlighting conserved and covarying nucleotides (color/marking per legend). Bottom: sequence logo (WebLogo) showing conservation within the SnoA seed region and the adjacent HcpR recognition motif in the target promoter. **B.** AlphaFold3-derived models of HcpR and SnoA. **C.** Modeled HcpR-SnoA-*hcp* promoter (P_hcp_) complex: left, overall view; right, magnified interfaces illustrating putative contacts between HcpR, SnoA, and promoter DNA. **D.** Predicted HcpR–SnoA–P_hcp_ architectures in representative *Bacteroides* species. **E.** Electrophoretic mobility shift assay: recombinant HcpR, promoter DNA (P_hcp_), and SnoA were incubated at 4 °C for 1 h and resolved on a native PAGE gel; DNA was visualized with SYBR Safe. **F.** As in (E), but SnoA was 5′-labeled with ATTO-680; fluorescence imaging highlights SnoA-containing complexes.

To test ternary complex formation experimentally, we incubated recombinant *B. thetaiotaomicron* HcpR with the *hcp* promoter DNA (P*_hcp_*) and/or SnoA and resolved complexes on native PAGE. Mixing HcpR, SnoA, and P*_hcp_*yielded a slowly migrating species with a higher apparent molecular weight than the HcpR-DNA complex, consistent with assembly of a ternary complex (**Fig. 4E**). This result was independent of detection method: covalent fluorescent labeling of SnoA produced a comparable shifted species that could be competed away by excess unlabeled SnoA, supporting binding specificity (**Fig. 4F**). Size-exclusion chromatography further showed that HcpR, SnoA, and P*_hcp_*co-elute with a profile indicative of complex formation (**Fig. S4F**). Together, these data support a model in which RNA arising from the SnoA locus can participate in an HcpR-containing promoter complex. This interaction may stabilize HcpR-promoter engagement or alter local promoter architecture to enhance transcriptional activation of nitrosative-stress defense genes.

### The SnoA locus promotes full activation of Hcp-dependent nitrosative defense

Having observed an interaction between RNA arising from the SnoA locus, the *hcp* promoter, and the transcription factor HcpR, we next sought to determine the contribution of the SnoA locus to the regulation of *B. thetaiotaomicron hcp* and *nrfA.* As a first step, we sought to uncouple *snoA* from *hcp* transcription. To do this, we generated a non-polar deletion of the *snoA*-*hcp* locus and complemented the mutant with either the wild-type sequence or a modified version in which the putative −10 and −30 elements^42^ of the *snoA* promoter were mutated to minimize promoter strength, while preserving the essential promoter elements of *hcp* and maintaining the native *hcp* coding sequence (**Fig. 5A**). qPCR analysis revealed that diminished *snoA* transcription (**Fig. S5A**) led to a marked reduction in *hcp* transcript levels during nitrosative stress (**Fig. 5B**). Supporting the notion that this effect is not simply due to polar interference to the *hcp* promoter, *nrfA* transcript levels were also significantly reduced, despite the *nrfA* locus being both (∼0.89 Mb away) distant and entirely intact (**Fig. 5C**). Notably, the wild-type strain outcompeted the *snoA* promoter mutant strain upon challenge with sodium nitrite, but not upon mock treatment (**Fig. 5D**). Conversely, increasing SnoA-locus dosage by inserting an additional copy at a neutral genomic site significantly increased *hcp* induction during low-dose nitrite challenge, whereas a sequence-scrambled control did not (**Fig. 5E**). Consistent with HcpR dependence, inserting an additional copy of the SnoA locus into the Δ*hcpR* strain did not significantly increase hcp induction (**Fig. S5B**).

**Fig. 5.**
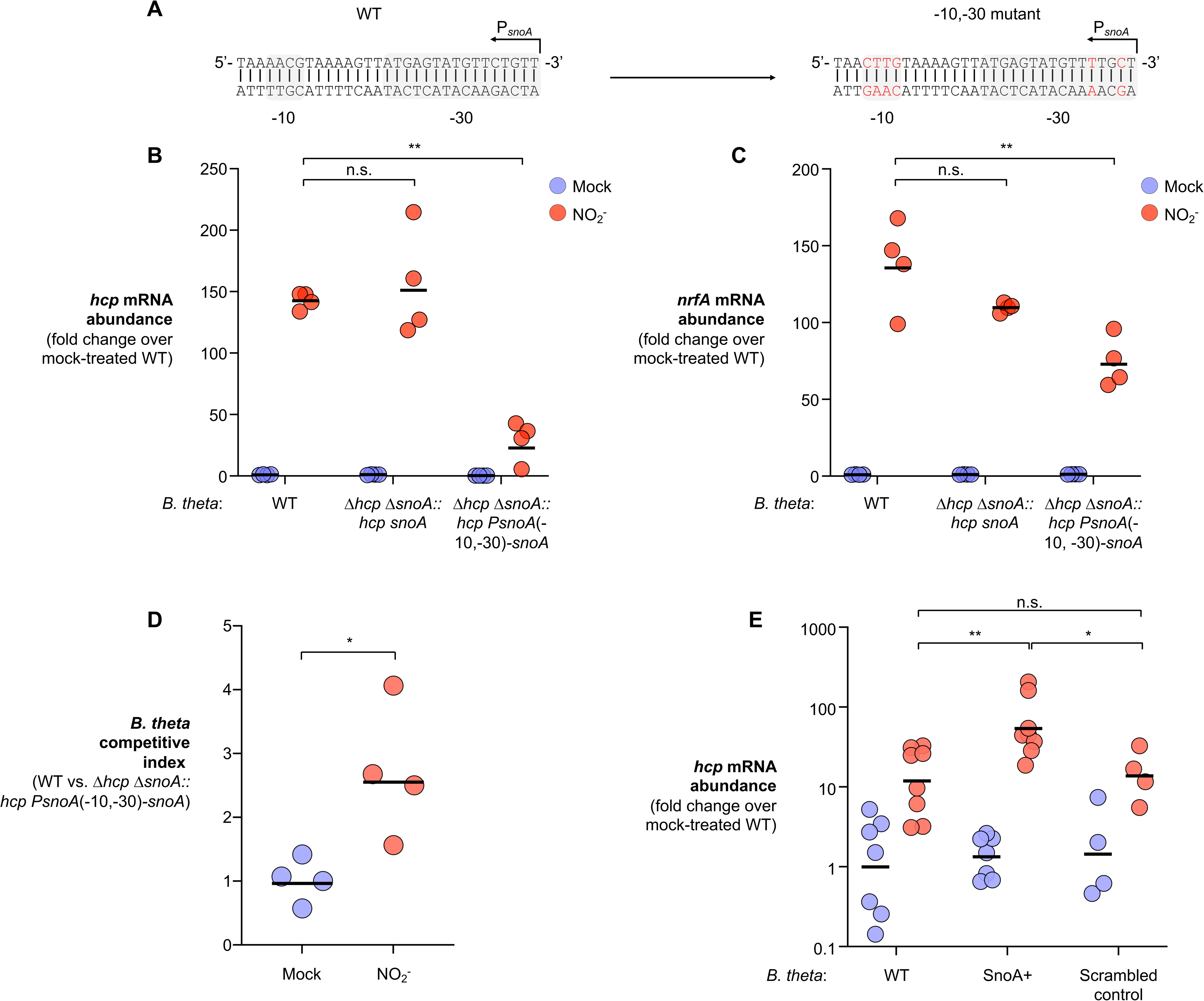
The SnoA locus promotes HcpR-dependent activation of the nitrosative-stress response. **A**. Schematic of the wild-type and mutant *snoA* promoter regions. Putative −10 and −30 promoter elements are indicated, with substituted nucleotides shown in red. **B-C**. RT-qPCR analysis of *hcp* (**B**) and *nrfA* (**C**) mRNA abundance in the indicated *B. thetaiotaomicron* strains following mock or nitrite treatment. Expression is shown as fold change relative to mock-treated WT. **D**. Competitive index of WT *B. thetaiotaomicron* versus the *snoA* promoter-mutant (τι*hcp* τι*snoA*::*hcp PsnoA*(−10, −30)-*snoA*) following mock or nitrite treatment. **E**. RT-qPCR analysis of *hcp* mRNA abundance in WT, SnoA+, or scrambled-control strains following mock or nitrite treatment. Expression is shown as fold change relative to mock-treated WT. Bars show geometric means. n.s., not significant; *, *P* < 0.05; **, *P* < 0.01.

### The HcpR-SnoA-Hcp axis promotes commensal resilience during host-derived intestinal nitrosative stress

The unique mechanism by which the NR-Hcp locus orchestrates nitrosative stress responses prompted us to test whether it sustains *B. thetaiotaomicron* resilience during gut inflammation. Because conventionally raised mice do not permit efficient *B. thetaiotaomicron* colonization^57^, we first treated C57BL/6 mice with an antibiotic cocktail to promote *B. thetaiotaomicron* engraftment^58^. We then colonized antibiotic-pretreated C57BL/6 mice with a 1:1 mixture of wild-type *B. thetaiotaomicron* and a nitrosative stress-defense mutant (ΔNR Δ*hcp*), followed by challenge by *Salmonella enterica serovar* Typhimurium (*S*. Tm), which activates robust mucosal immune responses through its Type III secretion systems and drives NO and nitrite production^59–62^ (**Fig. 6A-B, S6A**). Consistent with this, *S*. Tm infection induced marked mucosal inflammation and elevated luminal nitrite compared with mock-infected controls (**Fig. 6C-D, S6B-C**). In line with the idea that accumulated RNS threaten gut commensals, wild-type *B. thetaiotaomicron* significantly outcompeted the ΔNR Δ*hcp* mutant to a higher degree in both colon and cecum of *S*. Tm-infected mice than in sham-treated controls (**Fig. 6E&F**).

**Fig. 6.**
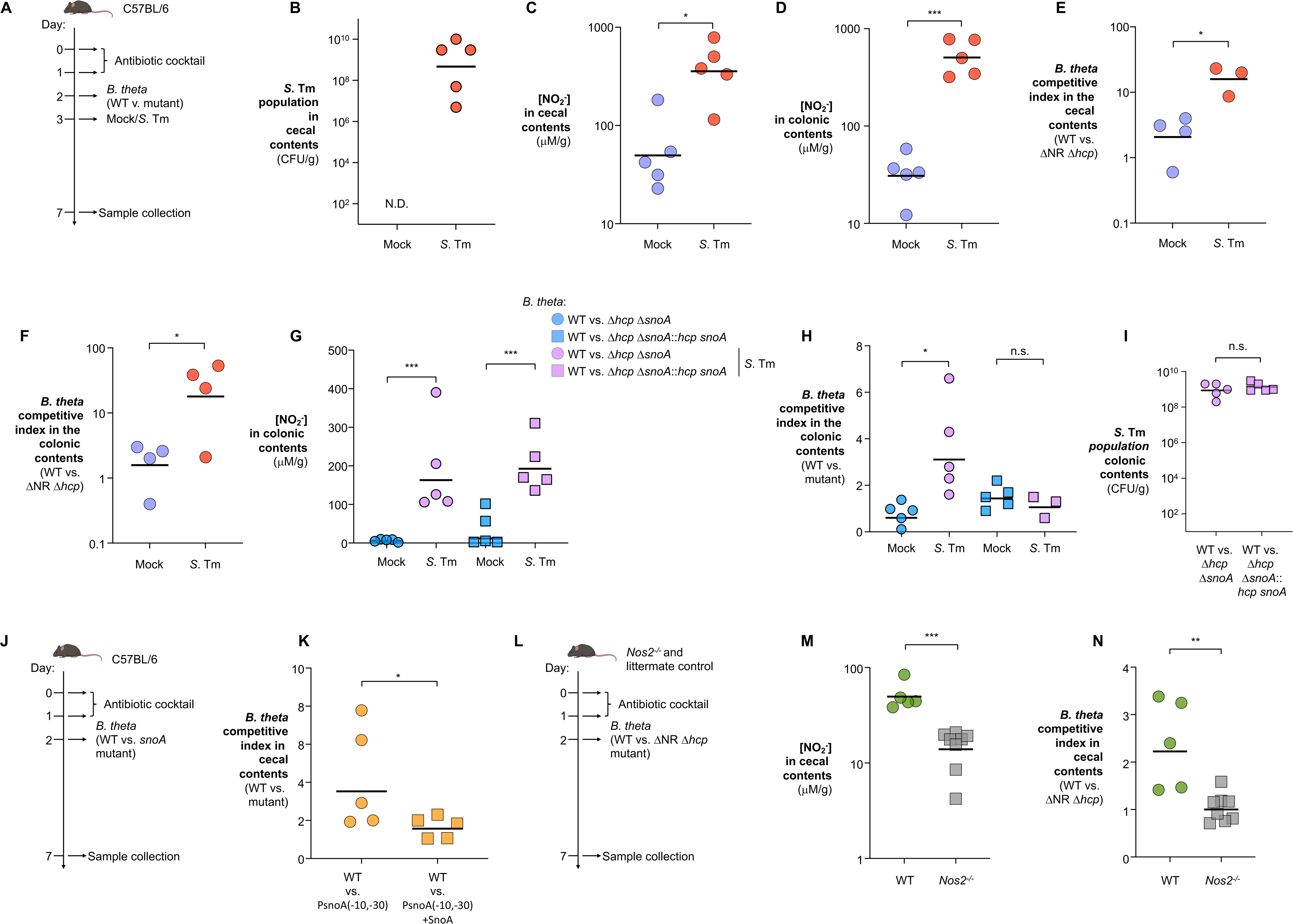
The HcpR-SnoA-Hcp axis promotes commensal resilience during host-derived intestinal nitrosative stress. **A-I** (*S*. Tm-infection model) Antibiotic-pretreated C57BL/6 mice were colonized with a 1:1 mix of *B. thetaiotaomicron* wild type and the indicated isogenic mutant(s). One day later, mice were mock-treated or challenged with *S.* Typhimurium (schematic in **A**). At 4 days post-infection (dpi), colonic and cecal contents were collected to measure luminal *S.* Typhimurium burden (**B, I**), nitrite quantified by Griess assay (**C-D, G**), and competitive index of WT over mutant by selective plating (**E-F, H**). **J-K,** antibiotic-perturbed infection-independent model. Antibiotic-pretreated C57BL/6 mice were colonized with a 1:1 mixture of WT and the *snoA* promoter mutant [P*_snoA_*(−10,−30)] or the promoter mutant carrying an additional SnoA copy in trans [P*_snoA_*(−10,−30) + SnoA] (schematic in **J**). **K**, competitive index of WT over the indicated mutant in cecal contents. **L-N**, Nos2-dependent infection-independent model. Antibiotic-pretreated *Nos2*^−/−^ mice and littermate controls were colonized with a 1:1 mixture of WT and the ΔNR Δ*hcp* mutant strain (schematic in **L**). M, cecal nitrite levels. **N**, competitive index of WT over ΔNR Δ*hcp* in cecal contents. Bars show geometric means (points = biological replicates). n.s., not significant; *, *P* < 0.05; **, *P* < 0.01; ***, *P* < 0.001.

We next asked how the SnoA-Hcp axis contributes to commensal fitness under nitrosative stress *in vivo*. Wild-type *B. thetaiotaomicron* was competed against either a Δ*hcp* Δ*snoA* mutant or a complemented strain (Δ*hcp* Δ*snoA*::*hcp snoA*) in *S*. Tm infected or mock-treated mice. As expected, *S*. Tm infection increased luminal nitrite (**Fig. 6G**). Under these conditions, wild-type *B. thetaiotaomicron* markedly outcompeted the Δ*hcp* Δ*snoA* mutant but not the complemented strain, whereas all strains exhibited comparable fitness in mock-treated animals (**Fig. 6H**). Differences in *B. thetaiotaomicron* fitness were not attributable to altered *S*. Tm burden or mucosal inflammation (**Fig. 6I, S6D**).

We next asked whether this pathway also protects *B. thetaiotaomicron* in an infection-independent inflammatory setting. Antibiotic treatment is commonly used to facilitate experimental colonization, but it can also perturb the microbiota and increase intestinal inflammatory tone, creating a host-derived stress environment even in the absence of an enteric pathogen^63^. We therefore used antibiotic-perturbed mice as a model of low-grade, infection-independent intestinal nitrosative stress.

In this model, wild-type *B. thetaiotaomicron* outcompeted the *snoA* promoter mutant [P*_snoA_*(−10,−30)], which exhibits markedly reduced *snoA* expression (**Fig. S5A**), whereas complementation with a wild-type copy of *snoA* at a neutral genomic site significantly rescued this fitness defect (**Fig. 6J,K**). These findings indicate that the SnoA locus promotes commensal fitness during antibiotic-associated intestinal stress.

We next asked whether this infection-independent fitness advantage depended on host-derived nitric oxide. To test this, groups of antibiotic-treated inducible nitric oxide synthase (*Nos2*)-deficient animals^64^ and littermate controls were colonized with a 1:1 mixture of wild-type *B. thetaiotaomicron* and the ΔNR Δ*hcp* mutant strain (**Fig. 6L)**. As expected, the *Nos2*-deficient mice displayed significantly lower cecal nitrite levels compared to littermate controls (**Fig. 6M)**. The two groups of mice differed significantly in their inflammatory cytokine expression only for *Nos2* (**Fig. S6E**). Consistent with our prior observation, the wild-type *B. thetaiotaomicron* strain exhibited a significant fitness advantage over the ΔNR Δ*hcp* mutant strain in the littermate control mice. However, this advantage was ablated in the *Nos2*-deficient mice (**Fig. 6N**). Together, these results indicate that the SnoA–Hcp/NR pathway contributes to B. thetaiotaomicron resilience during both infection-independent, antibiotic-associated nitrosative stress and enteric infection.

## Discussion

Host-derived reactive nitrogen species (RNS) are among the most potent chemical stresses encountered by gut microbes during inflammation^1^. Although pathogen adaptations to nitric oxide and nitrite have been dissected extensively, far less is known about how health-associated commensals withstand these same stresses. Here, we uncover a nitrosative-defense program in *B. thetaiotaomicron* that incorporates a mode of transcriptional regulation not previously appreciated: an RNA-associated transcriptional regulatory mechanism involving the SnoA locus and HcpR. These findings refine our understanding of how gut commensals maintain fitness during inflammatory episodes.

Our chemoproteomic mapping reveals that nitrosative stress broadly impacts the reactivity of cysteine residues in enzymes involved in central metabolism, amino acid and nucleotide biosynthesis, and numerous transport pathways in *B. thetaiotaomicron* (**Fig. 1**). These pathways mirror known vulnerabilities described in enteric pathogens^6^, underscoring the potential for shared biochemical liabilities that both pathogens and commensals face when exposed to NO and nitrite. However, *B. thetaiotaomicron* exhibits notable tolerance to RNS compared to model pathogens^65–67^, except under iron-limiting conditions—a context that reflects the inflammatory convergence of nitrosative and metal-starvation stresses. This tolerance likely arises from the complementary activities of NrfA and the hybrid cluster protein Hcp, which together enable detoxification of nitrite and NO and sustain metabolic continuity under stress. Yet their dependence on iron renders these systems an Achilles’ heel, offering a potential explanation for why commensals are depleted during mucosal inflammation, even when individual stressors appear tolerable in isolation. During iron limitation alone, *B. thetaiotaomicron* has been previously shown to downregulate expression of the nitrite reductase operon as part of its iron-sparing response *in vitro*^68^. However, in this study, iron limitation did not significantly reduce the nitrite-driven induction of *nrfA* expression, whereas the induction of *hcp* was almost completely ablated by iron limitation (**Fig. S4B**). Thus, nitrite reductase may serve as the prioritized nitrosative stress defense system when iron is limited. This is consistent with our observation that iron limitation is necessary for the *B. thetaiotaomicron* wild-type strain to outcompete the ΔNR mutant strain during nitrosative stress *in vitro* (**Fig. S2I**). Thus, the need to buffer simultaneous nitrosative and nutritional stresses provides a framework for understanding inflammation-driven dysbiosis^69,70^.

A second major finding of this study is that full activation of the Hcp nitrosative-stress defense pathway depends on the SnoA regulatory RNA locus. Unlike canonical post-transcriptional regulation by bacterial non-coding RNAs, the SnoA-associated RNA appears to act at the level of promoter-proximal transcriptional control. Genetic perturbation of the SnoA locus reduces *hcp* and *nrfA* induction, while biochemical assays show that SnoA-associated RNA can assemble with HcpR and promoter DNA *in vitro*. These findings support a model in which RNA arising from the SnoA locus enhances HcpR-dependent transcription, although the precise mature RNA species and the structural basis of activation remain unresolved.

Several lines of evidence support this model: (i) strong complementarity between SnoA and the −7 region of the *hcp* and *nrfA* promoters; (ii) HcpR-SnoA-DNA ternary complex formation *in vitro*; and (iii) separable contributions of HcpR and SnoA to promoter engagement.

From an evolutionary perspective, RNA-associated promoter regulation may provide *Bacteroides* species with expanded regulatory flexibility beyond palindromic or dyad-symmetric DNA motifs^71^. This added layer of specificity could enable rapid diversification of stress-response circuits. Indeed, comparative sequence analyses suggest that related HcpR-SnoA modules are present in distinct *Bacteroides* clades (**Fig. 4**), suggesting that this regulatory logic may extend beyond B. thetaiotaomicron while remaining clade-restricted. Our *in vivo* colonization assays demonstrate the physiological relevance of this system: *B. thetaiotaomicron* strains lacking SnoA and Hcp exhibit marked fitness defects only in the context of nitrosative inflammation, highlighting that this axis contributes to commensal persistence in the inflamed intestine.

This work raises several key questions. How widespread is RNA-associated promoter regulation among anaerobic gut commensals? Given the large, poorly characterized sRNA pools in *Bacteroidetes*, analogous mechanisms may regulate responses to other inflammatory stresses, such as antimicrobial peptides or nutrient limitation. What structural features enable HcpR to recognize DNA:RNA hybrids, and is this behavior unique to HcpR or shared among Crp-family regulators? Future cryo-EM structures should illuminate how RNA-associated promoter complexes are accommodated. Moreover, the upstream cues that induce SnoA transcription remain unresolved. Although heme binding supports a role for HcpR as a nitrosative sensor, HcpR is not required for SnoA induction, suggesting additional layers of redox or Fe-S stress sensing feed into this pathway. Finally, could RNA-associated promoter regulation be leveraged to stabilize beneficial microbes in conditions characterized by inflammation-driven dysbiosis, such as IBD or infectious colitis?

In summary, *B. thetaiotaomicron* mounts a multifaceted defense against nitrosative stress that includes an RNA-associated mechanism of promoter regulation. These findings deepen our understanding of how commensal microbes persist in the inflamed gut. As inflammation-driven perturbations of the microbiome are increasingly implicated in human disease, uncovering such hidden layers of stress adaptation will be essential for understanding and ultimately modulating microbiome resilience.

### Limitations of the Study

Several limitations should be acknowledged. First, the precise molecular identity of the SnoA-associated RNA remains unresolved. Although RNA-seq, RT-qPCR, promoter mutagenesis, complementation, and dosage experiments support nitrite-responsive transcription and regulatory function from this locus, we have not yet defined the mature transcript boundaries, processing state, or stoichiometry of the RNA species responsible for regulation. We therefore interpret SnoA as a nitrite-responsive regulatory RNA locus rather than a fully characterized canonical sRNA. Second, although biochemical assays support assembly of an HcpR-RNA-DNA complex, we have not resolved its high-resolution structure or directly defined how this complex affects RNA polymerase recruitment or transcription initiation. Finally, we have not assessed the long-term stability, off-target transcriptional consequences, or fitness trade-offs associated with altered SnoA-locus dosage in complex gut environments.

## Materials and Methods

### Bacterial strains, growth conditions, and *in vitro* competition assay

The bacterial strains used in this study are listed in the key resources table. *B. thetaiotaomicron* strains were grown anaerobically (90% N_2_, 5% CO_2_, 5% H_2_; vinyl anaerobic chamber, Coy Lab) in brain-heart-infusion supplemented (BHIS) media (0.8% brain heart infusion from solid, 0.5% peptic digest of animal tissue, 1.6% pancreatic digest of casein, 0.5% sodium chloride, 0.2% glucose, 0.25% disodium hydrogen phosphate, 0.005% haemin, 0.0001% vitamin K, pH 7.4) or BHIS plates (BHIS broth, 15 g/L agar) containing 50 μg/mL gentamicin (Gen) and 15 μg/mL chloramphenicol (Cm) or 25 μg/mL erythromycin (Erm) for 2 days at 37 °C. *E. coli* strains were routinely cultured in LB broth (10 g/L tryptone, 5 g/L yeast extract, 10 g/L sodium chloride) or on LB plates (LB broth, 15 g/L agar) at 37 °C supplemented with 100 µg/mL carbenicillin.

For the kinetic growth curves described in Figures 1 and 3, single colonies of the indicated strains were cultured anaerobically for 16 hours in BHIS. The strains were then inoculated into a 96-well plate at 1×10^5^ CFU/mL in BHIS supplemented with or without 100 µM bathophenanthroline disulfonic acid (BPS) and/or the indicated concentrations of nitric oxide (DPTA-NONOate), sodium nitrite, or sodium nitrate. Plates were incubated anaerobically at 37°C in a BioTek Epoch 2 plate reader, with a 1-minute linear shake and OD_600_ measurement every hour.

For *in vitro* competition assays, wild-type and mutant strains of *B. thetaiotaomicron* were cultured overnight as described above. Cultures were mixed at a 1:1 ratio and co-inoculated in BHIS, supplemented with 100 µM BPS, 50 µM DPTA NONOate (equivalent to 100 µM NO), and/or 100 µM sodium nitrite, as indicated. After 24 hours of incubation, cultures were serially diluted and plated onto selective BHIS agar containing the appropriate antibiotics. Plates were incubated anaerobically at 37 °C for 48 h, after which colony-forming units (CFUs) for each strain were enumerated.

### Plasmids

All the primers and plasmids used in this study are listed in the Key Resource Table. Suicide plasmids were routinely propagated in *E. coli* S17 λ*pir*. The flanking regions of *B. thetaiotaomicron* BT_1416-1418, BT_0687, BTnc284, and BT_0688 were amplified and assembled into pExchange-*tdk* using the Gibson Assembly Cloning Kit (New England Biolab, Boston) to give rise to pRF327, pRF328, pRF330, and pRF377 respectively.

### Construction of mutants by allelic exchange

All bacterial mutant strains constructed using the method below are listed in the Key Resource Table. For *B. thetaiotaomicron* mutants, suicide plasmid pExchange containing the flanking regions of genes of interest was conjugated using *E. coli* S17-1 λ*pir* as the conjugative donor strain into the *B. thetaiotaomicron*. Exconjugants with suicide plasmid integrated into the recipient chromosome were selected on BHIS plates supplemented with appropriate antibiotics. 5-fluoro-2-deoxy-uridine (FudR, 200 µg/mL in BHIS) plates were used to select for the second crossover event.

### Quantitative Cysteine Chemoproteomics

Cultures of wild-type *B. thetaiotaomicron* were cultured overnight as described above, before subculturing 1:50 into 50 mL BHIS. After 5 hours anaerobic incubation, the cultures were treated with a mock control, 1 mM sodium nitrite, or 0.5 mM PAPA NONOate. 1-hour post-treatment, the cells were pelleted at 3,200 xg and resuspended in 2.5 mL pre-reduced PBS supplemented with 1X Halt protease inhibitor cocktail. 0.5 mL of each sample was saved to quantify the total proteome. The remaining 2 mL of resuspended culture were lysed anaerobically via sonication at 40% amplitude, 10s on/10s off, for 3.5 minutes. The lysates were cleared at 13,200 rpm, 10 minutes, 4°C, and the protein concentration of the supernatant was quantified via Bradford assay for normalization to 2 mg/mL. Samples were assayed for both cysteine reactivity and protein abundance changes in pairwise chemoproteomic experiments comparing mock versus RNS-exposed (NO or nitrite) lysates.

Cysteine reactivity was quantified using isotopic activity-based protein profiling (isoTOP-ABPP)^35^. Briefly, mock and RNS-exposed proteomes (2 mg) were labeled with isotopic alkyne-functionalized iodoacetamide probes^72^ (100 μM; mock–iodoacetamide light (IAL); RNS-exposed – iodoacetamide heavy (IAH)) for 1 h at room temperature in the dark. Probe-labeled proteins were conjugated to photocleavable (PC) biotin-azide capture reagents via Cu(I)-catalyzed azide-alkyne cycloaddition (CuAAC; 100 µM PC biotin-azide; 1 mM TCEP; 100 µM TBTA; and 2 mM CuSO_4_) for 1 h at room temperature in the dark^73^. Next, light and heavy samples were precipitate (6,500 × g, 10 min, 4 °C), combined, washed with methanol, and resolubilized in SDS-containing PBS buffer. Samples were diluted to 0.2 % SDS, enriched on streptavidin resin, and extensively washed to remove non-specific interactions. On-bead reduction (10 mM DTT, 20 min, 65 °C) and alkylation (20 mM iodoacetamide, 30 min, 37 °C) were performed prior to overnight trypsin digestion (1 µg trypsin; 2 M urea in PBS, 1 mM CaCl_2_, overnight, 37 °C). Probe-labeled peptides were released by photocleavage of the linker, acidified, and desalted for LC–MS/MS analysis.

For complementary quantification of protein abundance, whole-proteome samples were analyzed by reductive dimethylation (ReDiMe)^74^. Proteins were precipitated (5% TCA), acetone washed, and resuspended in urea buffer (8 M urea in 100 mM TEAB), followed by reduction (15 mM DTT, 15 min, 75 °C), alkylation (12.5 mM iodoacetamide, 30 min, 37 °C), and tryptic digestion (1 µg trypsin; 1 M urea in 100 mM TEAB, 1 mM CaCl_2_, overnight, 37 °C). Peptides from anaerobic and oxygen-exposed samples were isotopically labeled with either light (mock) or heavy (RNS-exposed) formaldehyde (1% H_3_^12^CO or D_2_H^13^CO, 50 mM sodium cyanoborohydride, 2 h, room temperature), quenched with 1% ammonium hydroxide, combined, desalted, and fractionated by off-line high-pH reversed-phase chromatography^75^ prior to LC-MS/MS.

Peptides were analyzed by nanoLC–MS/MS on Orbitrap Exploris 240 mass spectrometer coupled to a Dionex Ultimate 3000 RSLCnano system in data-dependent acquisition (DDA) mode. MS1 scans (400-1800 MW, 120,000 resolution, RF lens 65%, AGC target 300%, automatic maximum injection time) were acquired every two seconds with dynamic exclusion enabled (repeat count 2, duration 10 s), followed by a variable number of data-dependent MS2 fragmentation scans (15,000 resolution, AGC 75%, maximum injection time 100 ms) of the most abundant precursor ions. MS2 analysis consisted of the isolation of precursor ions (isolation window 2 m/z), filtered for monoisotopic peak determination, theoretical isotopic envelope fit, intensity (5E4), and charge state (+2 – +6), followed by higher-energy collision dissociation (HCD, collision energy 30%). Spectra were searched using Thermo Proteome Discoverer V2.4 software package against a *B. thetaiotaomicron* UniProt database (UP000001414) using SequestHT and Percolator algorithms with trypsin specified as the protease (max of 2 missed cleavages) and precursor mass tolerance set to 10 ppm with a fragment mass tolerance of 0.02 Da. In addition to dynamic modifications representing oxidation of methionine (+15.995), as well as acetylation (+42.011) and/or methionine-loss (+131.040) of the protein N-terminus, appropriate static and variable modifications were set for cysteine alkylation and isotopic labeling. For ReDiMe analysis, static modifications on lysine and the peptide N-terminus of either +28.0313 (light) or +34.0632 (heavy) were specified, as well as a static modification on cysteine of +57.021 (alkylation). For isoTOP-ABPP, dynamic modifications on cysteine of +57.021 (alkylation) and either +285.159 (IAL) or +291.179 (IAH) were specified. Peptide and protein identifications were filtered to a false discovery rate ≤1%.

Relative cysteine reactivity and protein abundance were quantified from peptide light/heavy (ReDiMe) or heavy/light (isoTOP-ABPP) isotopic ratios using established pipelines (Proteome Discoverer)^35,76^. Protein-level ratios were calculated as the median of corresponding peptide measurements. All experiments were performed as two technical replicates of independent biological replicates (n = 2-3). Functional annotation of cysteine residues and protein localization was obtained from UniProt and integrated with quantitative datasets to identify RNS-sensitive cysteine sites across the proteome.

### Nitrosative stress RNA-seq

To identify nitrosative stress-responsive transcripts, 8 colonies of wild-type *B. thetaiotaomicron* were cultured anaerobically in 5 ml of brain-heart-infusion supplemented (BHIS) media (0.8% brain heart infusion from solid, 0.5% peptic digest of animal tissue, 1.6% pancreatic digest of casein, 0.5% sodium chloride, 0.2% glucose, 0.25% disodium hydrogen phosphate, 0.005% hemin, 0.0001% vitamin K, pH 7.4) for 24 hours, and subcultured (1:25) in 5 ml of BHIS for an additional 2.5 hours, until cultures reached early exponential phase. Half of the cultures were then treated with 1 mM sodium nitrite or mock control, and cultures were incubated for an additional 2 hours. Cell pellets were collected in RNAprotect (Qiagen), and RNA was extracted using TRI reagent (Molecular Research Center, Cincinnati). DNA contamination was removed using the Turbo DNA-free Kit (Ambion, USA) per the manufacturer’s recommendations.

To capture RNAs that originally had 5’ triphosphate, 5’ monophosphate, 5’ diphosphate or 5’ OH ends + 3’ OH or 3’ phosphate ends, RNA was dephosphorylated via treatment with Quick CIP (NEB), followed by treatment with T4 PNK (NEB), per the manufacturer’s instructions. RNA samples were then purified with the miRNeasy Micro Kit (Qiagen). Libraries were prepared using the RNAtag-seq method and sequenced on an Illumina Novaseq-SP platform (100 bp paired-end run). The sequencing reads generated during the current study are available at the European Bioinformatics Institute repository under accession No. ENA: PRJEB116001 (secondary accession ENA: ERP196150).

### Expression and purification of HcpR

Recombinant HcpR from *B. thetaiotaomicron* (222 aa; predicted MW 25.7 kDa, pI 8.9) was expressed in *E. coli* BL21(DE3) cells transformed with a pPL19 plasmid encoding N-terminally 10×His-tagged HcpR under control of the lac promoter. Cells were grown in LB medium supplemented with ampicillin (50 μg/mL) for a starter culture, which was used to inoculate autoinduction medium (Studier, 2005). Cultures were incubated at 37 °C for 4.5 h, followed by incubation at 25 °C for 18 h with shaking.

Cells were harvested by centrifugation and resuspended in lysis buffer (50 mM Tris-HCl pH 8.0, 1 M NaCl, 5% glycerol, 0.5 mM DTT) supplemented with EDTA-free protease inhibitors, PMSF, lysozyme, and DNase I. Cells were lysed by sonication and clarified by centrifugation at 4 °C for 60 min.

The supernatant was applied to Ni-NTA agarose pre-equilibrated in lysis buffer containing 20 mM imidazole. After washing with increasing imidazole concentrations (20–60 mM), bound protein was eluted with 300 mM imidazole. Eluted fractions were analyzed by SDS–PAGE, pooled, and dialyzed overnight at 4 °C against Buffer A (50 mM HEPES pH 7.0, 150 mM NaCl, 5% glycerol, 1 mM DTT, 1 mM EDTA).

Following dialysis, the sample was clarified by centrifugation and further purified by cation-exchange chromatography using a Mono S column equilibrated in Buffer A. Protein was eluted with a linear gradient of 150 mM to 1 M NaCl, and fractions were analyzed by SDS–PAGE. Peak fractions were pooled and subjected to size-exclusion chromatography on a Superdex 200 Increase column equilibrated in SEC buffer (20 mM HEPES pH 7.0, 150 mM NaCl, 5% glycerol, 1 mM DTT or TCEP, 1 mM EDTA). Purified HcpR was concentrated, flash-frozen, and stored at −80 °C.

### Hemin-Agarose Binding Assay

100 µL aliquots of hemin-agarose slurry (Sigma Aldrich) or control agarose slurry were washed 3 times in 1 mL binding buffer (50 mM Tris-HCl, 150 mM NaCl, pH 7.4). After washing, agarose pellets were resuspended in 1 mL binding buffer with 75 µg purified HcpR. The agarose and protein were incubated at 4°C for 1 hour, gently rocking. After 1 hour incubation, the samples were washed 3 times with 1mL binding buffer. Protein was then eluted by incubating agarose samples with 50 µL binding buffer supplemented with indicated conditions. Remaining bound protein was then eluted by boiling in Laemmli buffer. Input and sample eluents were resolved in an SDS-PAGE gel, and HcpR was visualized via staining with Coomassie Brilliant Blue R-250.

### Electrophoretic Mobility Shift Assay

A 270-bp DNA fragment containing the promoter region of *hcp* was amplified by PCR. Labeled SnoA was *in vitro* transcribed using HiScribe T7 High Yield RNA Kit (NEB) and aminoallyl-UTP-ATTO-680 (Jena Bioscience). Indicated combinations of the promoter, RNA, and purified HcpR were incubated for 1 hour at 4° in 0.5X TBE supplemented with 5% glycerol and 0.1 mg/mL BSA. After incubation, samples were resolved on a native TBE-PAGE gel at 4°C. DNA was visualized via 15-minute gel incubation in 1:10,000 SYBR Safe (Invitrogen) stain. Labeled RNA was visualized via fluorescence at 700 nm.

### Animal Experiments

All experiments were conducted in accordance with the policies of the Institutional Animal Care and Use Committee at Vanderbilt University Medical Center. C57BL/6J wild-type (cat# 000664), originally obtained from Jackson Laboratory (Bar Harbor), were procured, bred, and housed in sterile cages under specific pathogen-free conditions on a 12-hour light cycle, with *ad libitum* access to irradiated food and sterile water at Vanderbilt University Medical Center.

Seven to nine-week-old male and female mice were semi-randomly assigned into treatment groups before the experiment. Antibiotic cocktails (5 mg of each ampicillin (Sigma-Aldrich), metronidazole (Sigma-Aldrich), vancomycin (Chem Impex International), and neomycin (Sigma-Aldrich) per mouse) or mock treatment (water) were administered by oral gavage daily for 2 days. After antibiotic treatment, fecal pellets were collected and tested for bacterial growth on blood agar and blood agar supplemented with 50 μg/mL gentamicin. Only mice with no detectable bacterial growth on both media were included in the study to allow for quantification of experimentally-introduced *B. thetaiotaomicron* strains in luminal content and feces. At day 3, mice were inoculated with an equal mixture of 0.5 × 10^9^ CFU of the *B. thetaiotaomicron* wild-type strain and 0.5 × 10^9^ CFU of the indicated mutants. Two days later, mice were challenged by 1 × 10^9^ CFU of the *S.* Typhimurium strain SL1344 for 3 days.

For all experiments, mice were humanely euthanized at the indicated time points. After euthanasia, cecal and colonic tissue was collected, flash-frozen and stored at −80 °C for subsequent mRNA analysis or fixed in 10% formalin for histopathological analysis. For culture-dependent quantification of bacterial abundance, colonic and cecal contents were harvested in sterile PBS, and the load of *B. thetaiotaomicron* and *S.* Typhimurium were quantified by plating serial-diluted intestinal contents on selective agar.

Luminal nitrite levels were quantified via the Griess assay. 100 µL of cleared supernatant from homogenized intestinal contents samples were combined with 100 µL of a 1:1 mix of 0.1% N-(1-Naphthyl)ethylenediamine dihydrochloride and 2% sulfanilamide. Samples were incubated at 37°C for 10 minutes, then OD_540_ was measured. Nitrite concentrations were calculated from OD_540_ via a sodium nitrite standard curve prepared fresh for each experiment.

### Targeted quantification of mRNA levels in intestinal tissue and contents

Colonic or cecal tissue was homogenized in a bead beater (Precellys 24 Touch, Bertin Technologies) and RNA was extracted using TRI reagent (Molecular Research Center, Cincinnati). Total RNA from the cecum or colon content was extracted using the RNeasy PowerFecal Pro Kit (QIAGEN) per manufacturer’s instructions. DNA contamination was removed using the Turbo DNA-free Kit (Ambion, USA) per the manufacturer’s recommendations. SuperScript VILO cDNA Synthesis Kit (Thermo Fisher, USA) was used to generate cDNA. Real-time PCR was performed using PowerUp SYBR Green Master Mix (Applied Biosystem, USA), data were acquired in a CFX maestro 2.3 (Bio-Rad, USA). Target gene transcription of each sample was normalized to the respective levels of *Gapdh* (mouse) or *gmk* (bacterial) mRNA.

### Quantification and statistical analysis

Unless noted otherwise, data analysis was performed in GraphPad Prism v11.0.0. Values of bacterial population sizes, competitive indices, and fold changes in mRNA levels, were normally distributed after transformation by the natural logarithm. A two-tailed Student’s *t*-test was used for ln-transformed data. Unless otherwise stated, ^∗^, *P* < 0.05; ^∗∗^, *P* < 0.01; ^∗∗∗^, *P* < 0.001; ns, not statistically significant. In all mouse experiments, *N* refers to the number of animals from which samples were taken. The outlined statistical details of experiments, including the exact value of *N*, definition of center, and dispersion can be found in the figures or figure legends. Shapiro-Wilk test was used to determine the normality of log-transformed data. Sample sizes (i.e. the number of animals per group) were not estimated *a priori* since effect sizes in our system cannot be predicted. Mice that were euthanized early due to health concerns were excluded from analysis.

## Acknowledgements

Work in W. Z.’s lab was funded by the NIH (1R35GM147470, 1R01DK134692, R21AI187749, R21AI199418), The Pew Charitable Trust (2023-A-26048), V Foundation (V2022-032), Colorectal Cancer Alliance (10065978), and The G. Harold & Leila Y. Mathers Charitable Foundation (MF-2207-03128). Work in E. W.’s lab was funded by the NIH (R35GM134964). Work in Q. Z.’s lab was funded by NIH (R01MH132918). Any opinions, findings, conclusions, or recommendations expressed in this material are those of the author(s) and do not necessarily reflect the views of the funding agencies. The funders had no role in study design, data collection, and interpretation, or the decision to submit the work for publication.

## Author contributions

R. T. F. and W.Z. conceived the study and designed the experiments. R. T. F. performed and analyzed all in vitro experiments. R. T. F., M. L.-B., and L. S. performed and analyzed all in vivo experiments. D. W. B. and E. W. performed and analyzed the chemoproteomics experiments. R. T. F., L. C., D. S. and Q. Z. purified recombinant HcpR. R. T. F. performed the electrophoretic mobility shift assays (EMSAs) with advice from J. K. C. M. performed the covariance and phylogenetic analyses of the SnoA-HcpR-Hcp regulatory axis. R. T. F. performed bacterial RNAseq, J. L. and W. Z. analyzed the data. W. Z. and R. T. F. wrote the manuscript, and all authors reviewed and approved the final version.

