## Supplemental Figures for "A nitrite-responsive regulatory RNA locus sustains commensal resilience against nitrosative stress"

Fig. S1

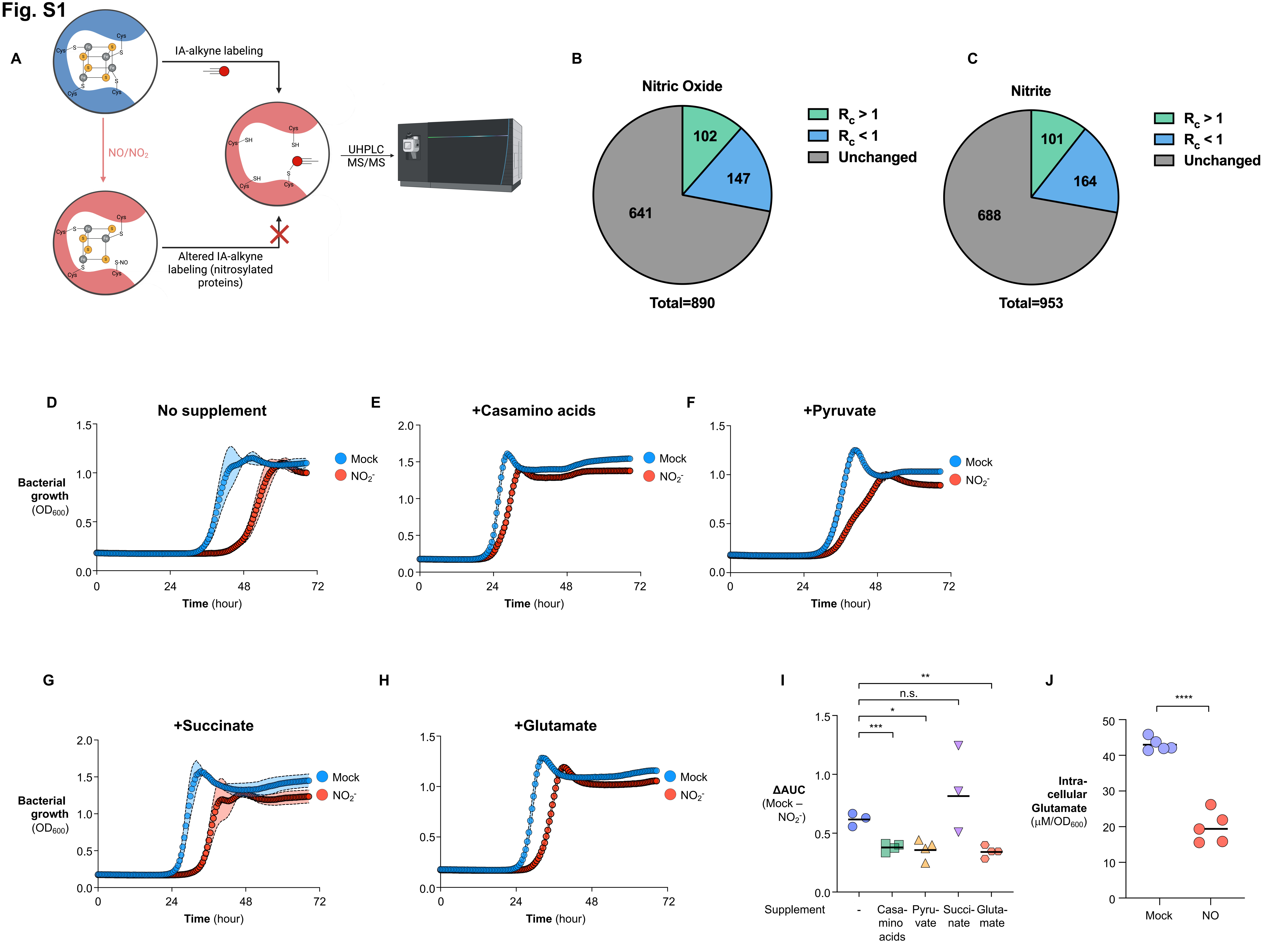

A
IA-alkyne labeling
NO/NO$_2$
UHPLC MS/MS
Altered IA-alkyne labeling (nitrosylated proteins)
Cys
SH
S-NO
B
Nitric Oxide
102
147
641
Total=890
$R_c > 1$
$R_c < 1$
Unchanged
C
Nitrite
101
164
688
Total=953
$R_c > 1$
$R_c < 1$
Unchanged
D
No supplement
Bacterial growth ($OD_{600}$)
Time (hour)
Mock
$NO_2^-$
E
+Casamino acids
Time (hour)
Mock
$NO_2^-$
F
+Pyruvate
Time (hour)
Mock
$NO_2^-$
G
+Succinate
Bacterial growth ($OD_{600}$)
Time (hour)
Mock
$NO_2^-$
H
+Glutamate
Time (hour)
Mock
$NO_2^-$
I
ΔAUC (Mock – $NO_2^-$)
Supplement
-
Casamino acids
Pyruvate
Succinate
Glutamate
n.s.
J
Intracellular Glutamate (μM/$OD_{600}$)
Mock
NO

**Fig. S1 (related to Fig. 1). Nitrosative stress impairs commensal fitness and remodels cysteine reactivity.**
**A.** Schematic of the IA-alkyne-based chemoproteomic workflow (isoTOP-ABPP) used to profile proteome-wide changes in cysteine reactivity after nitrosative stress. (**B,C**) Pie charts summarizing the extent of cysteine reactivity changes after exposure to NO (**B**) or nitrite (**C**). Cysteines with increased reactivity are shown as $R_C > 1$, cysteines with decreased reactivity are shown as $R_C < 1$, and unchanged cysteines are shown in gray. **D-H**. Growth of *B. thetaiotaomicron* in SDM challenged with 1 mM sodium nitrite with 10 mM of the indicated supplements: no supplement (**D**), casamino acids (**E**), sodium pyruvate (**F**), sodium succinate (**G**), glutamate (**H**). Bacterial growth ($OD_{600}$) was monitored over time. **I.** Difference in area under the curve (AUC) between mock-treated and nitrite-challenged SDM for *B. thetaiotaomicron* kinetic growth curves with indicated supplements. **J.** Subcultures of *B. thetaiotaomicron* were mock-treated or challenged with nitric oxide (PAPA NONOate) for 1 hour. Cultures were pelleted and washed with PBS. Metabolites were extracted using 80% methanol and glutamate quantified via LC/MS. Bars show geometric means (points = biological replicates). n.s., not significant; *, $P < 0.05$; **,$P < 0.01$; ***,$P < 0.001$.

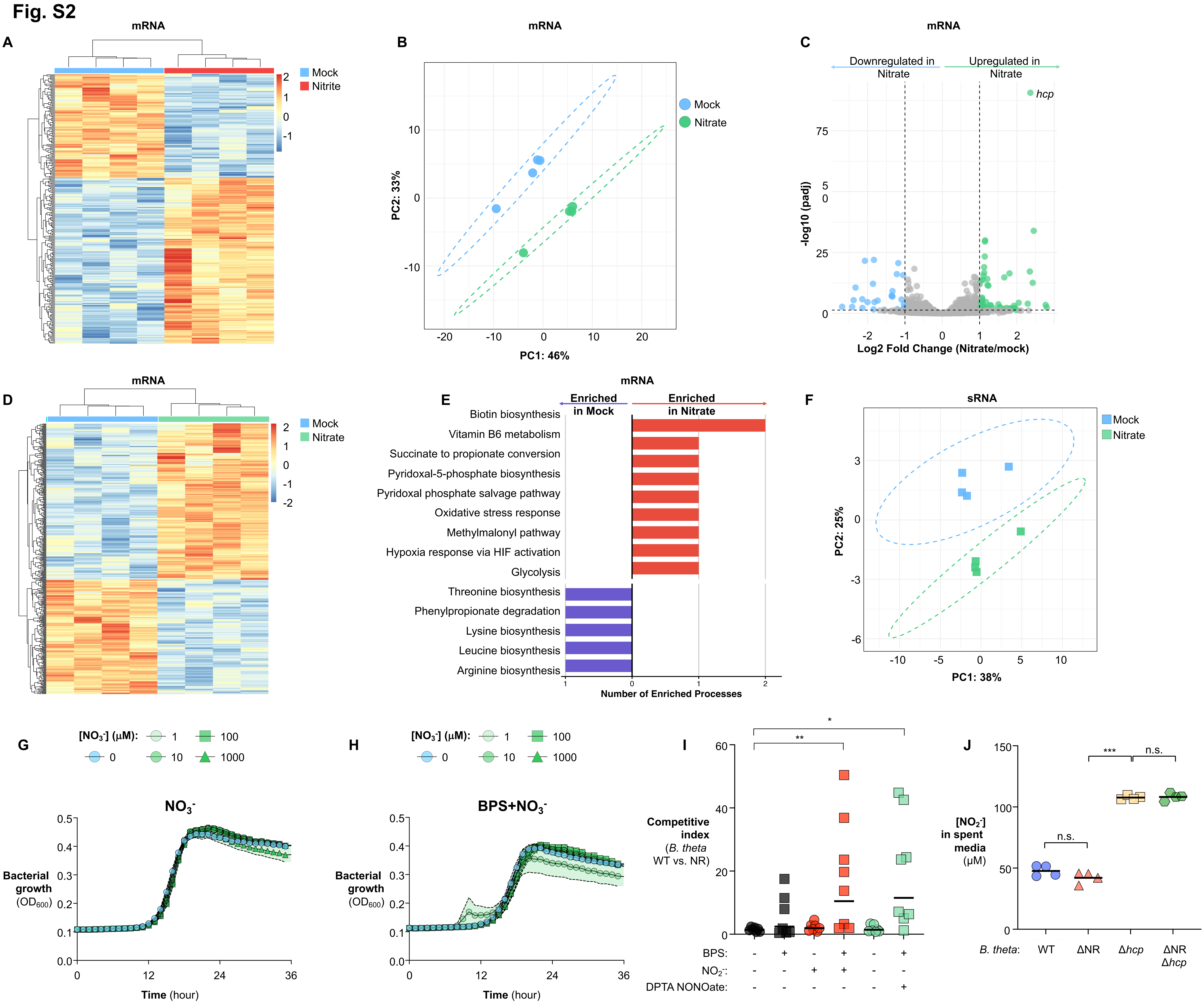
Fig. S2
A
mRNA
Mock
Nitrite
B
mRNA
Mock
Nitrate
PC1: 46%
PC2: 33%
C
mRNA
Downregulated in Nitrate
Upregulated in Nitrate
hcp
-log10 (padj)
Log2 Fold Change (Nitrate/mock)
D
mRNA
Mock
Nitrate
E
mRNA
Enriched in Mock
Enriched in Nitrate
Biotin biosynthesis
Vitamin B6 metabolism
Succinate to propionate conversion
Pyridoxal-5-phosphate biosynthesis
Pyridoxal phosphate salvage pathway
Oxidative stress response
Methylmalonyl pathway
Hypoxia response via HIF activation
Glycolysis
Threonine biosynthesis
Phenylpropionate degradation
Lysine biosynthesis
Leucine biosynthesis
Arginine biosynthesis
Number of Enriched Processes
F
sRNA
Mock
Nitrate
PC1: 38%
PC2: 25%
G
[NO3-] (μM):
NO3-
Bacterial growth (OD600)
Time (hour)
H
[NO3-] (μM):
BPS+NO3-
Bacterial growth (OD600)
Time (hour)
I
Competitive index (B. theta WT vs. NR)
BPS:
NO2-:
DPTA NONOate:
J
[NO2-] in spent media (μM)
n.s.
***
B. theta:
WT
ΔNR
Δhcp
ΔNR Δhcp

**Fig. S2 (related to Fig. 2). *B. thetaiotaomicron* mounts a protective transcriptional program to detoxify nitrosative stress.**

**A.** Heatmap of nitrite-altered mRNA transcriptome after 1 h nitrite exposure. **B-E.** Nitrate control (mRNA). **B.** principal component analysis (PCA); **C.** volcano plot; **D.** heatmap; **E.** PANTHER biological-process enrichment of differentially expressed mRNAs. **F.** Principal component analysis (PCA) of the nitrate control (sRNA). **G-I** Growth of *B. thetaiotaomicron* in BHI under iron-limiting conditions (BPS) with the indicated nitrosative stressors. Bacterial growth ($OD_{600}$) was monitored over time. **G.** nitrate under iron replete condition; **H.** nitrate under iron limited condition (BPS). **I.** A 1:1 mixture of *B. thetaiotaomicron* wild-type and an isogenic nitrite reductase (NR)-deficient mutant was inoculated in BHI with iron limitation (BPS) ± nitrosative stress ($NO_2^-$ or NO). Competitive index was determined by selective plating. **J.** Quantification of nitrite in spent BHI media 72 hours after indicated *B. thetaiotaomicron* strains at exponential phase were treated with 1 mM sodium nitrite. Bars (where shown) depict geometric means. n.s., not significant; *, $P < 0.05$; **, $P < 0.01$; ***, $P < 0.001$. BPS, bathophenanthroline disulfonate; $NO_2^-$, nitrite; $NO_3^-$, nitrate; DPTA NONOate, Dipropylenetriamine NONOate (nitric oxide producer).

Fig. S3

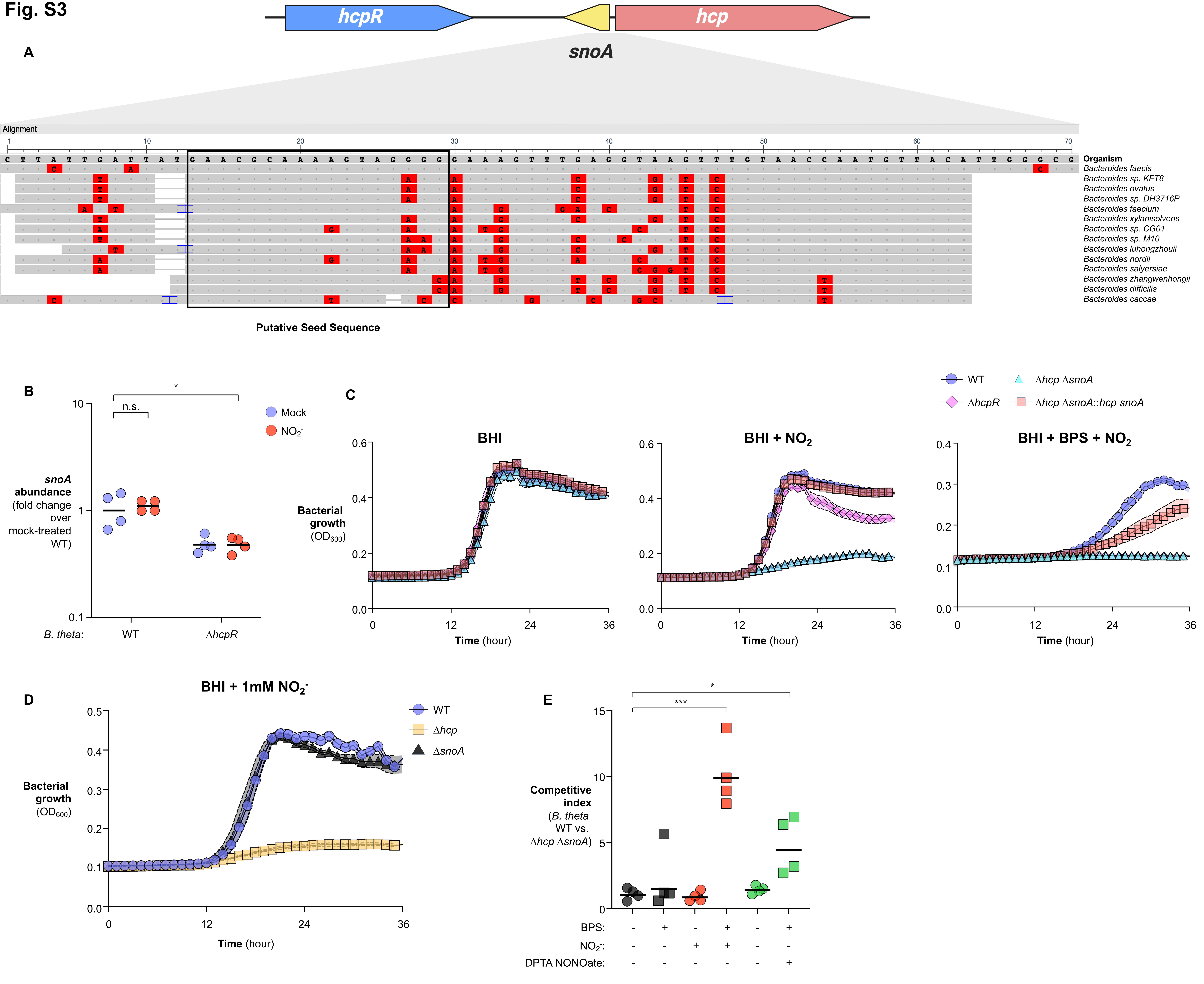

hcpR
hcp
A
snoA
Alignment
Organism
Bacteroides faecis
Bacteroides sp. KFT8
Bacteroides ovatus
Bacteroides sp. DH3716P
Bacteroides faecium
Bacteroides xylanisolvens
Bacteroides sp. CG01
Bacteroides sp. M10
Bacteroides luhongzhouii
Bacteroides nordii
Bacteroides salyersiae
Bacteroides zhangwenhongii
Bacteroides difficilis
Bacteroides caccae
Putative Seed Sequence
B
n.s.
*
Mock
NO2-
snoA abundance (fold change over mock-treated WT)
B. theta:
WT
ΔhcpR
C
WT
ΔhcpR
Δhcp ΔsnoA
Δhcp ΔsnoA::hcp snoA
BHI
BHI + NO2
BHI + BPS + NO2
Bacterial growth (OD600)
Time (hour)
D
BHI + 1mM NO2-
WT
Δhcp
ΔsnoA
Bacterial growth (OD600)
Time (hour)
E
***
*
Competitive index (B. theta WT vs. Δhcp ΔsnoA)
BPS:
NO2-:
DPTA NONOate:

**Fig. S3 (related to Fig. 3). The SnoA regulatory RNA locus modulates nitrosative-stress defense in *B. thetaiotaomicron*.**

**A.** Conservation of SnoA across bacteria (sequence alignment/phylogeny) predicted by CopraRNA, highlighting the preserved seed region and promoter-recognition features. **B**. SnoA abundance in the indicated strains after exposure to nitrite or mock control. **C-D**. Growth of *B. thetaiotaomicron* cultures ($OD_{600}$) under iron limitation with nitrosative stress. **E**. A 1:1 mixture of *B. thetaiotaomicron* wild-type and an isogenic *hcp snoA*-deficient mutant was inoculated in BHI with iron limitation (BPS) ± nitrosative stress ($NO_2^-$ or NO). Competitive index was determined by selective plating. Bars show geometric means. n.s., not significant; *, $P < 0.05$; ***, $P < 0.001$.

Fig. S4

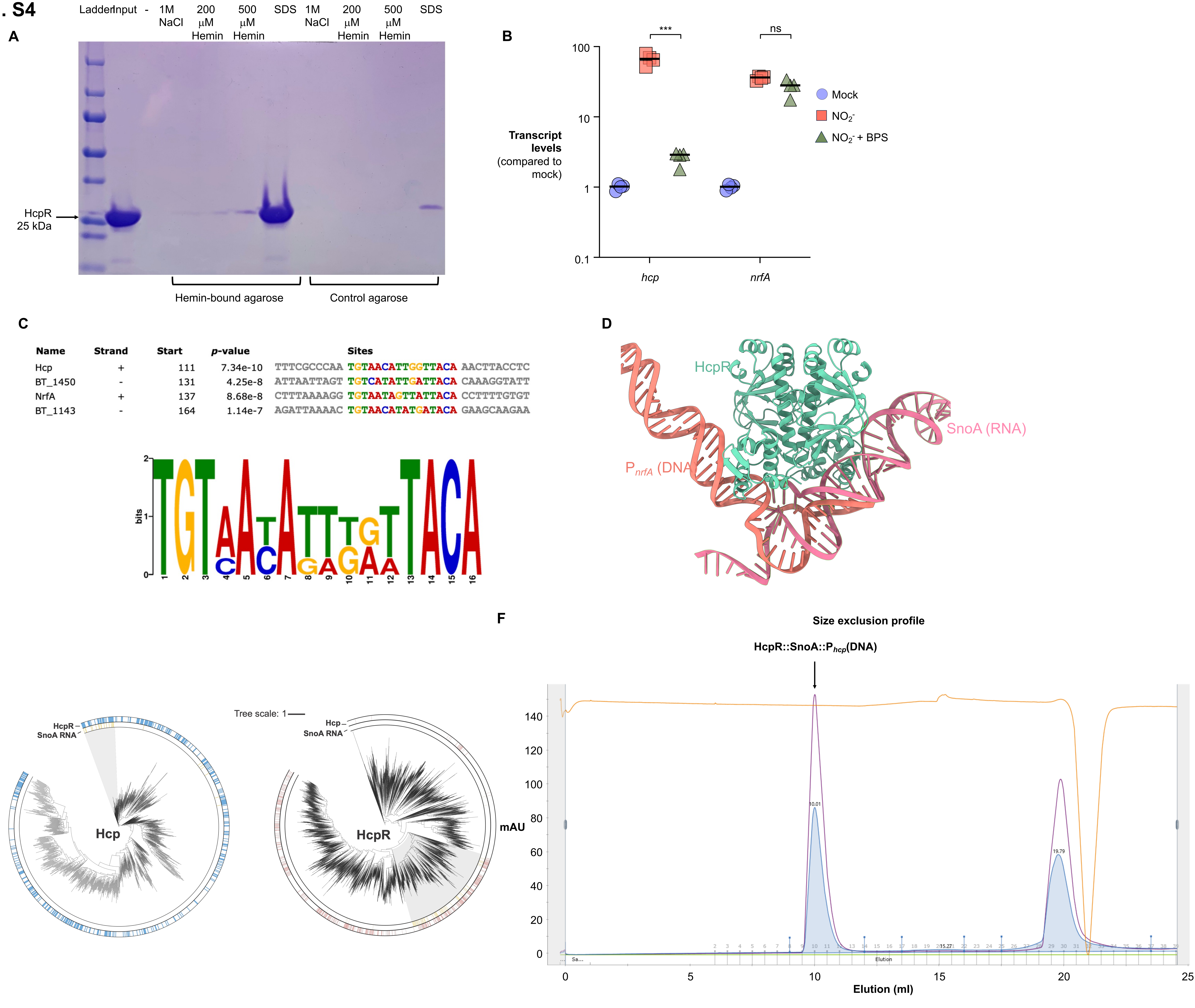
A
Ladder
Input
-
1M NaCl
200 µM Hemin
500 µM Hemin
SDS
1M NaCl
200 µM Hemin
500 µM Hemin
SDS
HcpR 25 kDa
Hemin-bound agarose
Control agarose
B
***
ns
Transcript levels (compared to mock)
100
10
1
0.1
hcp
nrfA
Mock
NO2-
NO2- + BPS
C
Name Strand Start p-value Sites
Hcp + 111 7.34e-10 TTTCGCCCAA TGTAACATTGGTTACA AACTTACCTC
BT_1450 - 131 4.25e-8 ATTAATTAGT TGTCATATTGATTACA CAAAGGTATT
NrfA + 137 8.68e-8 CTTTAAAAGG TGTAATAGTTATTACA CCTTTTGTGT
BT_1143 - 164 1.14e-7 AGATTAAAAC TGTAACATATGATACA GAAGCAAGAA
bits
D
HcpR
SnoA (RNA)
PnrfA (DNA)
E
HcpR
SnoA RNA
Hcp
Tree scale: 1
Hcp
SnoA RNA
HcpR
F
Size exclusion profile
HcpR::SnoA::Phcp(DNA)
mAU
10.01
19.79
Elution (ml)

**Fig. S4 (related to Fig. 4). SnoA-associated RNA forms HcpR-containing complexes with conserved promoter elements.**

**A.** HcpR binds to heme: HcpR was incubated with heme-conjugated agarose at 4 °C overnight, beads were washed, and bound protein was eluted with NaCl, increasing concentrations of free heme, or SDS. Fractions were resolved by SDS-PAGE. **B.** RT-qPCR of *hcp* and *nrfA* mRNA in wild-type *B. thetaiotaomicron* after exposure to mock control, nitrite, or nitrite and BPS. **C.** MEME analysis of promoters from top 4 upregulated genes in nitrite RNAseq identifies putative HcpR consensus sequence. **D.** AlphaFold3-predicted model of the HcpR-SnoA complex bound to the *nrfA* promoter ($P_{nrfA}$), illustrating the putative interfaces between protein, RNA, and DNA. **E.** Comparative phylogenetic analysis of the SnoA–HcpR–Hcp regulatory module. Midpoint-rooted circular phylogenies of Hcp (left) and HcpR (right) homologs. Gray shading highlights clades containing SnoA-associated RNAs identified by the covariance model. Outer-ring annotations indicate the presence of nearby HcpR proteins (left, blue), Hcp proteins (right, pink), and SnoA-associated RNAs (yellow) located within a ±5 kb genomic window. The phylogenies suggest that the SnoA–HcpR–Hcp regulatory architecture is conserved within distinct clades rather than universally across all homologs. **F.** Recombinant HcpR, promoter DNA ($P_{hcp}$), and SnoA were incubated at 4 °C for 1 h and subjected to size-exclusion chromatography on a Superdex 200 column to resolve complex formation. Elution profiles correspond to free components and assembled complexes. Bars show geometric means. n.s., not significant; ***, $P < 0.001$.

### Fig. S5

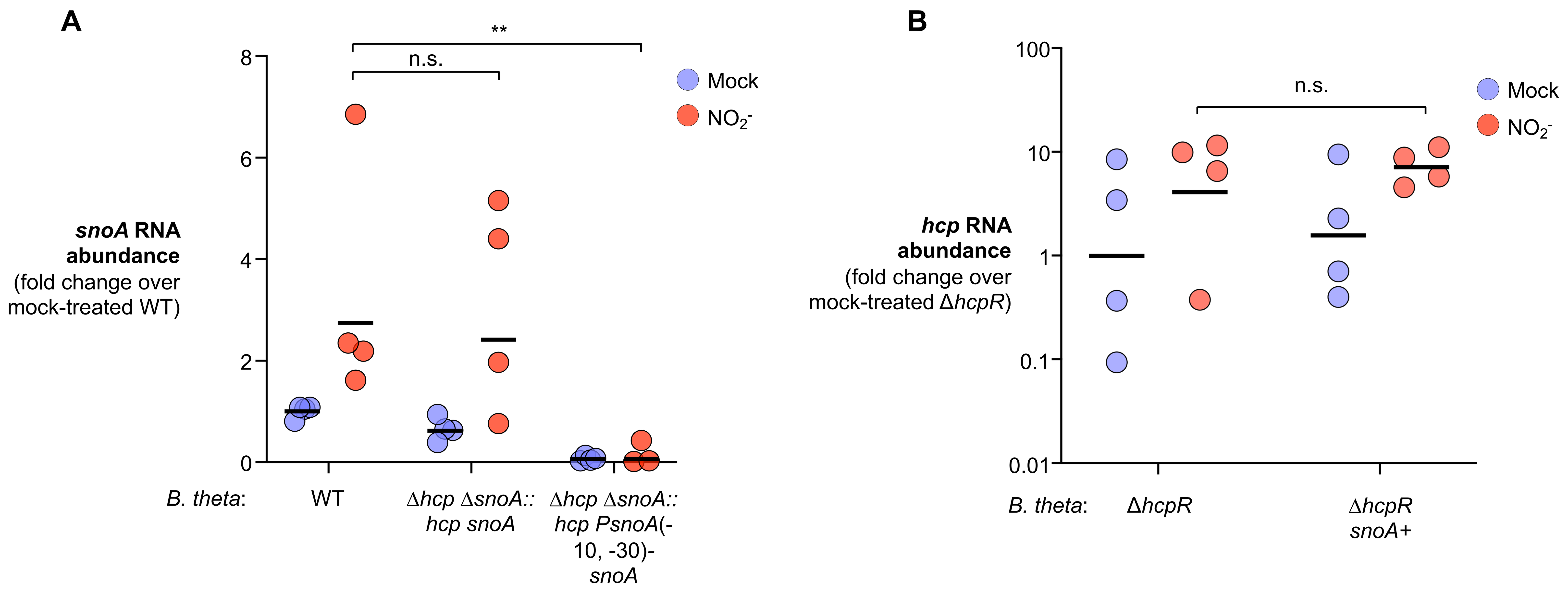
A
snoA RNA abundance (fold change over mock-treated WT)
8
6
4
2
0
n.s.
**
Mock
$NO_2^-$
B. theta:
WT
Δhcp ΔsnoA:: hcp snoA
Δhcp ΔsnoA:: hcp PsnoA(-10, -30)-snoA
B
hcp RNA abundance (fold change over mock-treated ΔhcpR)
100
10
1
0.1
0.01
n.s.
Mock
$NO_2^-$
B. theta:
ΔhcpR
ΔhcpR snoA+

**Fig. S5 (related to Fig. 5). The SnoA locus promotes HcpR-dependent activation of the nitrosative-stress response.**

**A**. RT-qPCR analysis of snoA RNA abundance in the indicated *B. thetaiotaomicron* strains following mock or nitrite treatment. Expression is shown as fold change relative to mock-treated WT. **B**. RT-qPCR analysis of hcp mRNA abundance in Δ*hcpR* or Δ*hcpR snoA*+ strains following mock or nitrite treatment. Expression is shown as fold change relative to mock-treated Δ*hcpR*. Bars show geometric means. n.s., not significant; **, $P < 0.01$.

### Fig. S6

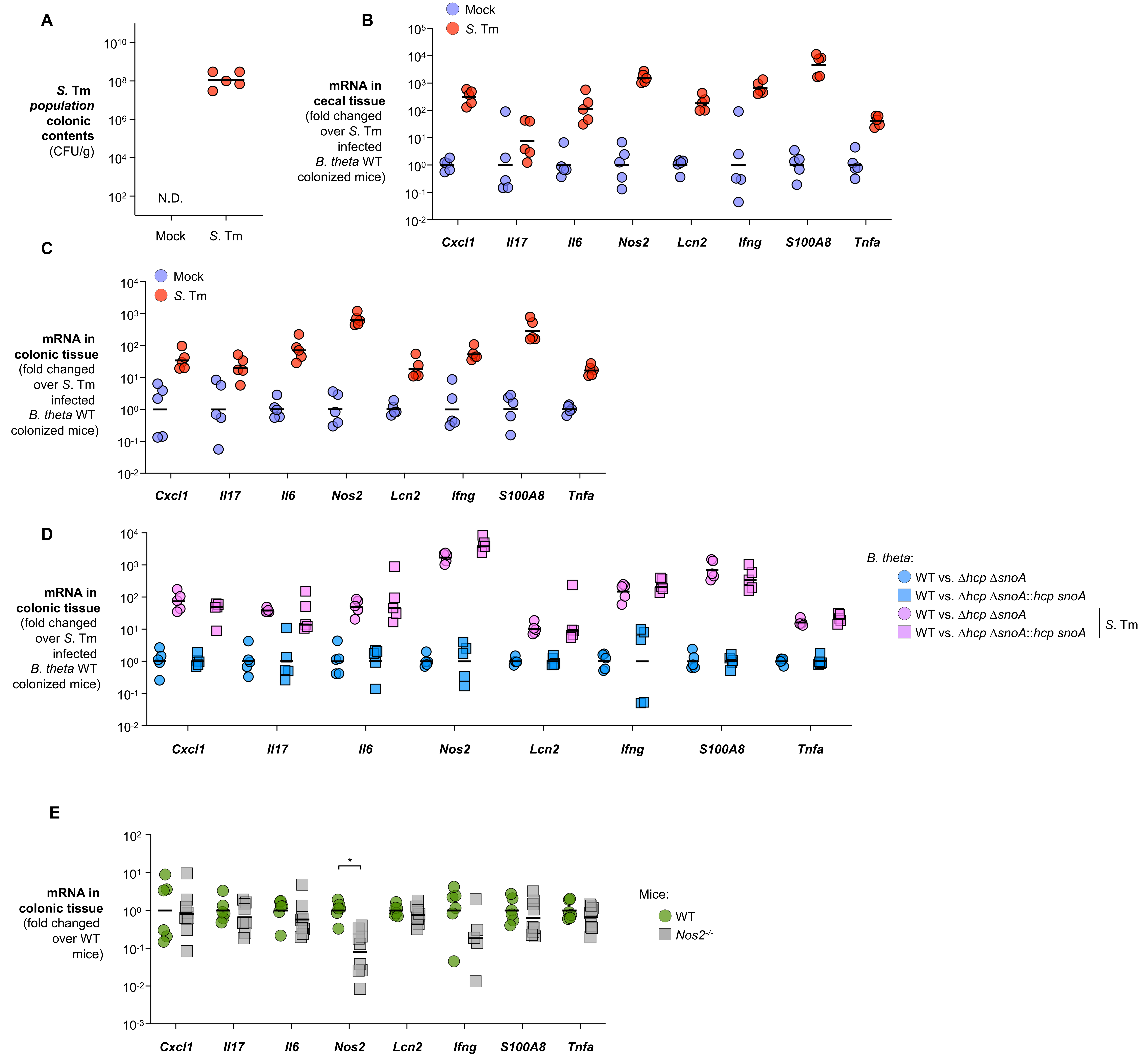

A
S. Tm population colonic contents (CFU/g)
N.D.
Mock
S. Tm
B
Mock
S. Tm
mRNA in cecal tissue (fold changed over S. Tm infected B. theta WT colonized mice)
Cxcl1
Il17
Il6
Nos2
Lcn2
Ifng
S100A8
Tnfa
C
mRNA in colonic tissue (fold changed over S. Tm infected B. theta WT colonized mice)
D
B. theta:
WT vs. Δhcp ΔsnoA
WT vs. Δhcp ΔsnoA::hcp snoA
S. Tm
E
mRNA in colonic tissue (fold changed over WT mice)
*
Mice:
WT
Nos2-/-

**Fig. S6 (related to Fig. 6). The HcpR-SnoA-Hcp axis promotes commensal resilience during host-derived intestinal nitrosative stress.**

**A-D** (*S*. Tm infection model). Antibiotic-pretreated C57BL/6 mice were colonized with a 1:1 mix of *B. thetaiotaomicron* wild type and the indicated isogenic mutant(s) or complemented strain. One day later, mice were mock-treated or challenged with *S.* Typhimurium. At 4 days post-infection (dpi), *S*. Tm population in the colonic contents was determined by selective plating (**A**), colonic and cecal tissue were collected and the mRNA levels of indicated cytokines were quantified using RT-qPCR (**B, C, D**). **E** (*Nos2*$^{-/-}$ uninfected model) Groups of antibiotic-pretreated C57BL/6 *Nos2*$^{-/-}$ mice and littermate controls were colonized with a 1:1 mix of the *B. thetaiotaomicron* WT and the ΔNR Δ*hcp* mutant strain. Cecal tissue was collected and cecal tissue inflammatory gene expression was measured by RT-qPCR and normalized to WT mice. Bars show geometric means. *, $P < 0.05$.
